# Retinoic Acid Inducible Gene-I like Receptors Activate Snail and Slug to Limit RNA Viral Infections

**DOI:** 10.1101/2020.10.27.357509

**Authors:** Dhiviya Vedagiri, Divya Gupta, Anurag Mishra, Gayathri Krishna, Meenakshi Bhaskar, Anirban Basu, Debasis Nayak, Manjula Kalia, Mohanan Valiya Veettil, Krishnan Harinivas Harshan

**Affiliations:** CSIR-Centre for Cellular and Molecular Biology, Uppal Road, Hyderabad-500007, India; Discipline of Biosciences and Biomedical Engineering, Indian Institute of Technology Indore, Simrol, Indore-453552, India; Virology Laboratory, Department of Biotechnology, Cochin University of Science and Technology, Cochin, Kerala-682022, India; National Brain Research Centre, Manesar, Haryana-122052, India; Regional Centre for Biotechnology, NCR Biotech Science Cluster, Faridabad, Haryana-121001, India; Academy for Scientific and Innovative Research (AcSIR), Ghaziabad-201002, India

**Keywords:** Epithelial-Mesenchymal Transition-Transcription Factors (EMT-TFs), Innate antiviral response, Interferon Regulatory Factor 3 (IRF3), Interferon Stimulated Response Element (ISRE), RNA virus

## Abstract

RLRs sense cytosolic non-self RNAs including viral RNAs before mounting a response leading to the activation of Type-I IFNs. Here, we identify a previously unknown regulation of Snail, a transcription regulator known in EMT, during RNA viral infections and describe its possible implication. RNA viral infections, poly (I:C) transfection and ectopic expression of RLR components activated Snail and Slug in epithelial cells. Detailed examination revealed that MAVS and phosphorylated IRF3 are essential in this regulation. We identified two ISREs in *SNAI1* promoter region and their alterations rendered the promoter non-responsive to phospho-IRF3 in luciferase assay. Ectopic expression of Snail and Slug activated RLR pathway and dramatically limited RNA viral infections in epithelial cells pointing to their antiviral functions. Thus, Snail and Slug are transcriptionally regulated by RLRs in a similar manner as IFN-β and they in turn promote RLR pathway possibly strengthening the antiviral state in the cell.

## INTRODUCTION

Epithelial cells are early barriers that encounter viral infections. Despite being immune generalists, they mount a robust innate antiviral immune response, thereby producing type-I interferons (IFN) that limit viral spread (1). Pathogen associated molecular patterns (PAMP) are detected by pattern recognition receptors (PRR), further assisting the infected cells in mounting responses against pathogens. Nucleic acids of viral origin are strong PAMPs that trigger a cellular response through cytosolic PRRs (2). RIG-I and MDA5 are two such PRRs that recognize RNA of virus origin (3–6). The two N-terminal CARD domains of these molecules allow them to oligomerize and to interact independently with the mitochondrial antiviral signalling protein (MAVS) that subsequently oligomerizes through its CARD-like domains on the outer membrane of mitochondria. In turn, these events recruit TBK1 and IRF3, leading to IRF3 phosphorylation on several residues in its C-terminal domain by TBK1 (7–9). Phosphorylated IRF3 undergoes homodimerization before translocating to the nucleus where it associates with promoter regions of Type I IFN genes, IFN-α and IFN-β, through ISRE elements. (7, 10)

Epithelial-mesenchymal transition (EMT) is a biological process with essential functions in embryo development, wound healing, and cancer metastasis (11–13). During EMT, epithelial cells leave true epithelialness to acquire varying degrees of mesenchymalness. A set of transcription factors from the Snail, Twist, and Zeb families, collectively referred to as EMT-TFs, facilitate the transition through major transcriptional reprogramming (14, 15). They suppress several epithelial markers, including junction proteins E-cadherin and Claudin, and activate mesenchymal markers such as Vimentin and Fibronectin (14). Some of the key signal pathways that regulate EMT-TFs are TGF-β, Wnt/β-Catenin, PI3K-AKT/GSK-3β, Jagged, Notch, Hedgehog, and Hippo (16). EMT provides enormous flexibility to the cellular phenotype and behavior, which are well exploited by the three events mentioned earlier.

Several recent reports have demonstrated EMT induction during infection by oncoviruses such as hepatitis B (HBV)(17), human papilloma (HPV)(18), Epstein Barr (EBV) (19), Kaposi’s sarcoma-associated herpesvirus (KSHV) and hepatitis C (HCV)(20) viruses, both *in vitro* and *in vivo.* Various viral proteins also activate key regulatory pathways activating EMT-TFs (21). One speculated potential outcome of this EMT is the progression of cancer induced by these viruses. Notwithstanding their contribution to cancer progression, viruses are unlikely to induce EMT at initial periods of infections with an impact on cancer that develops several years post-infection. Further, induction of EMT by a few non-oncoviruses such as Ebolavirus (EBOV)(22), Respiratory Syncytial Virus (RSV) (23), Human Cytomegalovirus (HCMV)(24), Human Rhinovirus (HRV) (25) and also HIV(26) point to the likelihood of its unidentified roles in viral infection. Nevertheless, there is a lacuna in our understanding of EMT activation by viruses from diverse families through common mechanisms and the biological consequence of such activation on the infection.

In this study, we addressed two significant questions on virus-induced EMT. First, we sought to identify a universal pathway used by viruses to induce EMT. Here, we identified that the RLR-IRF3 pathway that is employed to regulate Type-I IFN expression during viral infections also regulate Snail and Slug. In the second, we investigated the consequences of EMT on viral infections. This universal mechanism is likely used by all the cells in response to RNA viruses. We also establish that EMT-TFs have significant antiviral properties against RNA viruses. Through these studies, we identified a previously unknown mechanism of activation of Snail and Slug by innate antiviral pathways that, in turn, facilitates the sustenance of the antiviral response.

## RESULTS

### EMT is a universal response to RNA virus infection

In order to test the possibility of EMT being a typical response in viral infection, we investigated if it is induced in response to infections by common non-onco RNA viruses Dengue (DENV), Japanese encephalitis (JEV), and Vesicular stomatitis virus (VSV) in common epithelial cancer cell lines. A549 cells robustly express E-cadherin with modest levels of Vimentin, Snail, and Slug (Figure S1A). Huh7.5 cells express E-cadherin, Vimentin, and Snail, but no Slug. MCF-7 expresses robust levels of epithelial marker E-cadherin and undetectable levels of mesenchymal marker Vimentin. EMT-TFs Snail and Slug were also undetected in them. We noticed a significant drop in E-cadherin and increased Vimentin levels in A549 and Huh7.5 cells infected with DENV, indicating EMT (Figure 1A). A similar change in E-cadherin levels in MCF-7 cells was visible without affecting Vimentin levels. Similar to DENV, JEV infection also brought about changes in EMT markers (Figure 1A). In agreement, infected cells reported elevated levels of *SNAI1, SNAI2,* and *ZEB1* transcripts (Figure 1 B-D). Remarkable activation of *ZEB2* transcripts in A549 but not in MCF-7 and Huh7.5 cells upon JEV and DENV infection (Figure S1B) suggests cell-specific activation of EMT-TFs and their redundancy in viral infections. At transcript levels, lower *CDH1,* and higher *VIM* in infected samples demonstrated induction of EMT transcriptional reprogramming by these (+) stranded RNA viruses (Figure S1 C-E). Strong activation of *VIM* transcripts in MCF-7 cells shows transcriptional regulation consistent with EMT (Figure S1 E), but the absence of its induction at protein level points to additional post-transcriptional regulatory events. VSV, a (-) stranded RNA virus, induced *SNAI1* substantially, and *SNAI2* and *ZEB1* moderately (Figure 1E). The presence of GFP confirmed VSV infection in infected cells compared to mock-infected cells (Figure S1F). These results illustrate that RNA virus infections induce EMT as a general cellular response. Additionally, they indicate that EMT early during viral infections need not necessarily be intended at promoting oncogenesis.

**Figure 1.**
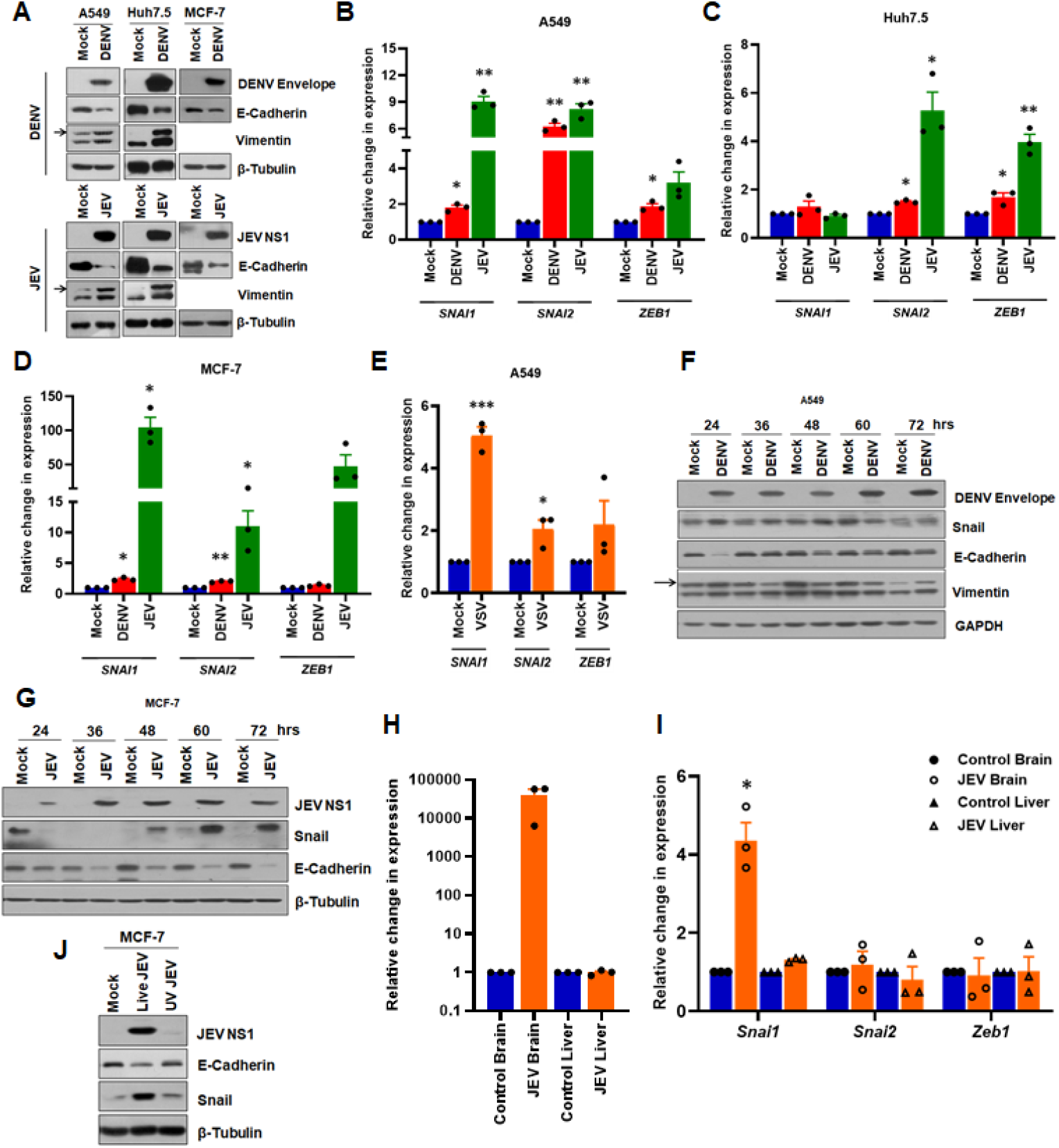
Non-oncoviruses induce EMT: (**A**) Immunoblots of DENV and JEV infected A549, Huh7.5, and MCF-7 cells. Cells were infected with either JEV at 0.1 MOI for 48 hrs or DENV at 0.5 MOI for 72 hrs. Mock and virus-infected cells were lysed, electrophoresed, and blotted for respective molecules. (**B** - **D**) Real-time qRT-PCR analysis of panel (**A**) samples, quantified for *SNAI1, SNAI2,* and *ZEB1* transcripts and normalized against *GAPDH.* (**E**) *SNAI1, SNAI2,* and *ZEB1* transcripts quantified in mock and VSV infected A549 cells by real-time qRT-PCR and normalized to *GAPDH.* (**F**) Time kinetics of DENV infected A549 cells immunoblotted for DENV envelope and EMT markers at indicated time points. Infection was performed as indicated in (A). (**G**) Time kinetics of EMT in MCF-7 infected with JEV for specified time points by immunoblotting. (**H** & **I**) Real-time PCR analysis of control and JEV infected mice tissues. *JEV NS3* levels (**H**) were quantified from the dissected brain and liver of control and JEV infected mice. Simultaneously, *Snai1, Snai2,* and *Zeb1* transcripts (**I**) were also quantified and normalized to *ActB.* (**J**) Immunoblot analysis of MCF-7 cells infected with either mock or live or UV-inactivated JEV for 48 hrs and probed for JEV NS1, E-Cadherin, Snail, and β-Tubulin.

To gain further insight into the interplay between virus infection and EMT markers, EMT was analyzed over a time course spanning from 24-72 hpi. Detection of DENV envelope from 24 hpi in infected A549 cells confirmed the infection (Figure 1F). Substantial E-Cadherin depletion and a concomitant increase in Vimentin were evident in the infected cells. Elevated Snail in the infected cells indicated the onset of EMT as early as 24 hpi. The activation of Snail and depletion of E-Cadherin was also evident in JEV infected MCF-7 cells (Figure 1G). A gradual activation of *SNAI1* and *SNAI2* transcripts was visible from 36 hpi (Figure S1G). Thus, our observations indicate that EMT is set early during infection and remains throughout the infection period tested.

We further tested if EMT is induced *in vivo.* Intraperitoneal injection of JEV in mice resulted in increased *Snai1* levels in the brain where high JEV RNA levels were present (Figure 1H), but not in the liver where JEV RNA was absent, demonstrating that viral infection could induce EMT-TFs *in vivo* (Figure 1I). Juxtaposing our results with the existing literature suggests that EMT induction is a general response to viral infection.

EMT could be induced as a consequence of the interaction between the virion and cell surface receptors, viral replication, or through signaling activities promoted by viral proteins. In order to test this, we infected MCF-7 cells with UV inactivated JEV. The inactivated virus could not establish infection, as is evident from Figure 1J. Interestingly, the inactivated virus also did not induce EMT, unlike the infectious virus suggesting that post-entry events are critical for EMT during viral infections.

### EMT-TFs Snail and Slug are induced by viral RNA

Several viral proteins having no shared features among them have been demonstrated to induce EMT (19, 27, 28). Since various RNA viruses across different classes induce EMT, we reasoned that viral RNA molecules could be behind it as they are sensed by PRRs sense it based on their common molecular patterns. This hypothesis was tested by transfecting MCF-7 cells with total RNA purified from JEV or mock-infected cells and analyzing EMT at multiple time intervals. RNA transfection was confirmed by the presence of JEV RNA at 12 h post-transfection (hpt) (Figure 2A) and by the robust activation of antiviral *IFNB1* and *IFIT1* transcripts (Figure 2B). Substantial *SNAI1* activation was seen in cells transfected with RNA from JEV infected cells compared with the control cells that received RNA from mock-infected cells (Figure 2C). The activation was visible from 12 hpt, intensified until 48 hpt, and remained at significant levels thereafter. *SNAI2* and *ZEB1* transcripts were also activated, albeit by lower magnitude from 36 hpt. Significant activation of *VIM* was evident from 36 hpt, while *CDH1* levels dropped by 60 hpt (Figure 2D). These results support the idea that viral RNA can induce EMT, despite the possibility that viral proteins generated from transfected RNA could also have contributed to the process.

**Figure 2:**
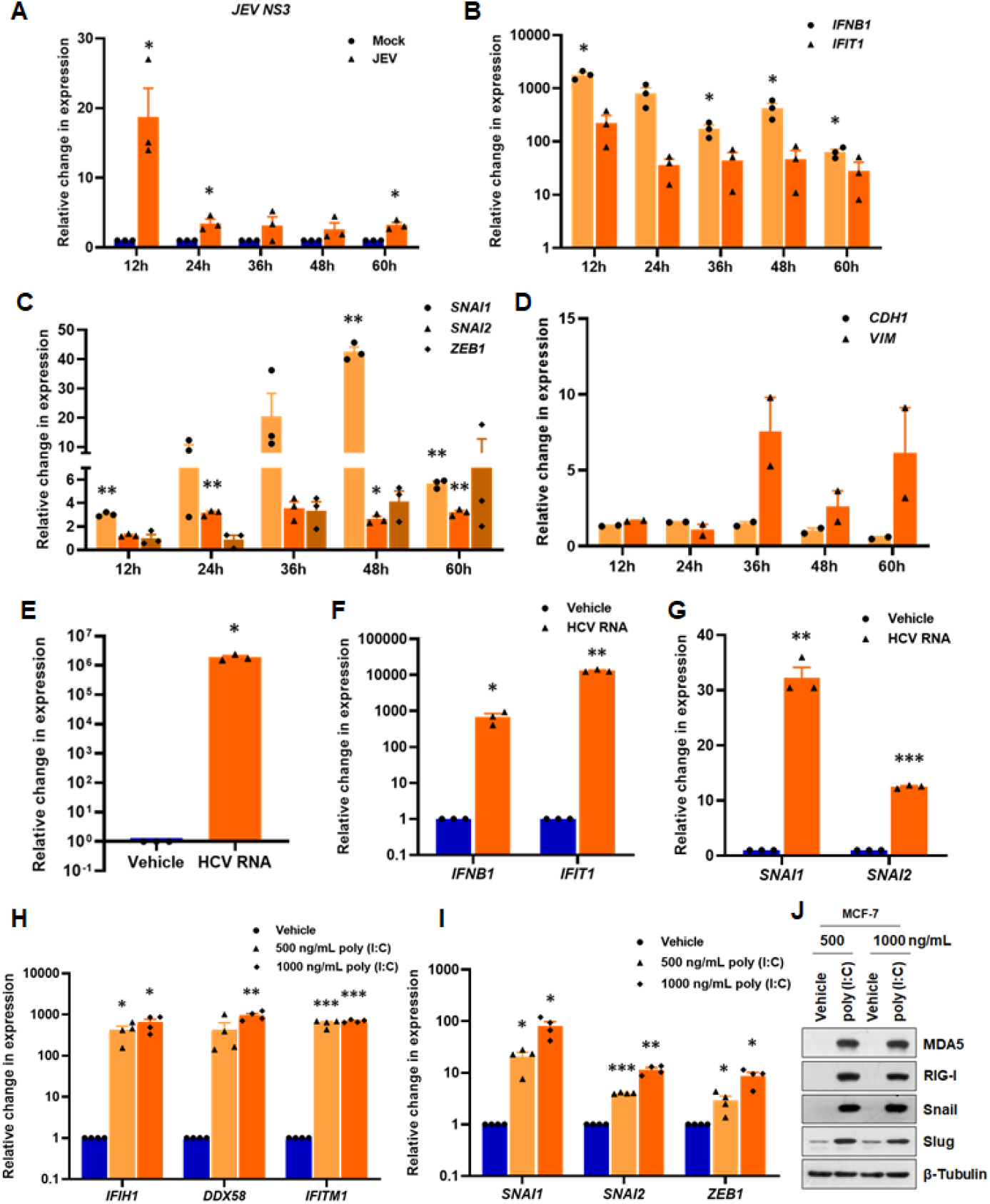
EMT-TFs are activated by viral RNA: (**A** - **D**) RNA prepared from mock or JEV infected MCF-7 was transfected into MCF-7 cells, and cells were harvested at indicated time points (mock and JEV cellular RNA pool designated as mock and JEV). RNA isolated from transfected cells was reverse transcribed and analyzed for (**A**) *JEV NS3* levels, (**B**) antiviral genes, *IFNB1* and *IFIT1,* (**C**) EMT-TFs, *SNAI1, SNAI2* and *ZEB1,* and (**D**) EMT markers, *CDH1* and *VIM* transcripts were quantified by real-time qRT-PCR. GAPDH was used as the normalization control. (E - G) Real-time qRT-PCR of MCF-7 transfected with purified IVT HCV RNA for 48 hrs. (**E**) *HCV RNA* was quantified from vehicle and IVT HCV RNA transfected MCF-7 cells. Simultaneously, innate immune genes, *IFNB1* and *IFIT1* (**F**), and EMT-TFs, *SNAI1* and *SNAI2* (**G**) were also quantified. All transcripts were normalized to *β-Tubulin* mRNA. (**H** - **J**) Poly (I:C) treatment of MCF-7 by transfection. (**H**) Antiviral genes, *IFIH1, DDX58,* and *IFITM1,* (**I**) EMT-TFs, *SNAI1, SNAI2,* and *ZEB1,* were quantified by real-time qRT-PCR, and relative change in expression with vehicle transfected cells was calculated. *GAPDH* was used as the normalization control. (**J**) Immunoblots of poly (I:C) transfected MCF-7 cells.

To validate this observation, we transfected *in-vitro* transcribed HCV ssRNA into MCF-7 cells. HCV RNA is not expected to replicate in non-hepatic MCF-7 cells, and hence the RNA would remain in ss-form. Among other potential factors, the absence of *miR-122,* a liver enriched factor key for HCV RNA stabilization and replication, could be a reason for this non-permissiveness of MCF-7 (29). HCV RNA transfection (Figure 2E) elicited a robust antiviral response in MCF-7 cells, as shown by elevated *IFNB1* and *IFIT1* transcripts (Figure 2F). Interestingly, HCV RNA substantially induced *SNAI1* and *SNAI2* in MCF-7 (Figure 2G). The activation of *SNAI1* and *SNAI2* in them suggests that single-stranded viral RNA can induce EMT-TFs just as it activates type-I IFN. These results indicated the involvement of the primary RNA sensors RIG-I and MDA5 that regulate type-I IFN production in response to RNA virus infection.

To further confirm the above findings, we performed transfection of dsRNA mimic poly (I:C) that is well known to activate the RLR pathway. Transfection of 500 ng/mL poly (I:C) into MCF-7 cells triggered significant antiviral response evidenced by robust activation of *IFIH1, DDX58* (encoding MDA-5 and RIG-I respectively) and *IFITM1* (Figure 2H). As seen in HCV RNA transfection, poly (I:C) transfection significantly activated *SNAI1* and lesser but still considerable extent *SNAI2* and *ZEB1* (Figure 2I). This finding was substantiated by immunoblotting (Figure 2J). Transfection of 1000ng/mL poly (I:C) further up-regulated the EMT-TF transcripts. *SNAI2* and *ZEB2* were robustly activated, while *SNAI1* activation was modest in poly (I:C) transfected A549 (Figure S2A and B) cells. Consistent activation of Slug with no change in Snail by the treatment in A549 indicated the redundancy displayed between the EMT-TFs across cell types (Figure S2C). Further strengthening this data, poly (I:C) transfection in HEK293 cells activated *SNAI1* and *SNAI2* transcripts while only Snail levels were elevated (Figure S2 D and E). These results confirm that RNA sensing pathways modulate the expression of various EMT-TFs based on the cell context and could be a potential mechanism of activation of EMT during RNA viral infections.

### RIG-I and MDA5 regulate EMT-TFs expression

Since poly (I:C) mounts antiviral response through dsRNA binding proteins RIG-I and MDA5, we tested if they could induce EMT-TFs expression. Over-expression of MDA5, RIG-I, and MAVS independently activated *SNAI1* and *SNAI2* but not *ZEB1* (Figure 3A) in MCF-7 cells while Snail showed activation at protein level (Figure 3B). Thus, it is apparent that these modulators of antiviral molecules transcriptionally activate the two major EMT-TFs in addition to mounting an antiviral response. Given that MAVS is crucial in assimilating upstream signals from the RLRs, we tested if its loss negatively impacts EMT-TFs. In agreement, MAVS KO A549 cells expressed less Snail (Figure 3C), while ectopic expression of MAVS elevated the Snail levels (Figures 3 D and E). We further tested if their activation is restricted upon poly (I:C) transfection, and interestingly, *SNAI2* and *ZEB2* activation were suboptimal in MAVS KO cells as compared with the WT-cells upon poly (I:C) transfection (Figure 3F). As expected, MAVS KO cells displayed inadequate antiviral response, as evident from lower activation of *IFIH1, DDX58, IFITM3,* and *IFNB1* (Figure S3A). JEV infection also caused limited activation of *SNAI1* in MAVS KO compared with the WT-cells (Figure 3G). Further, expression of CARD deletion mutant of MAVS (ΔCARD-MAVS) failed to induce Snail unlike the FL-MAVS (Figure 3 H and I). These results collectively demonstrated that dsRNA sensing machinery transcriptionally activates EMT-TFs Snail and Slug. The activation of Snail and Slug by dsRNA sensing machinery raised the vital question on the point of intervention in their regulation. MAVS coordinates TBK1/IKKε mediated phosphorylation of IRF3 that subsequently activates Type I IFN transcription. We tested if IFN signaling is critical to EMT-TF regulation by treating MCF-7 cells with IFNα-2a. While this treatment activated ISGs such as IFIT1 and IFITM1, it failed to induce EMT-TFs both at protein (Figure 3J) and transcript levels (Figure S3 B and C), suggesting that IFN production is dispensable for the regulation of EMT-TFs during viral infection.

**Figure 3:**
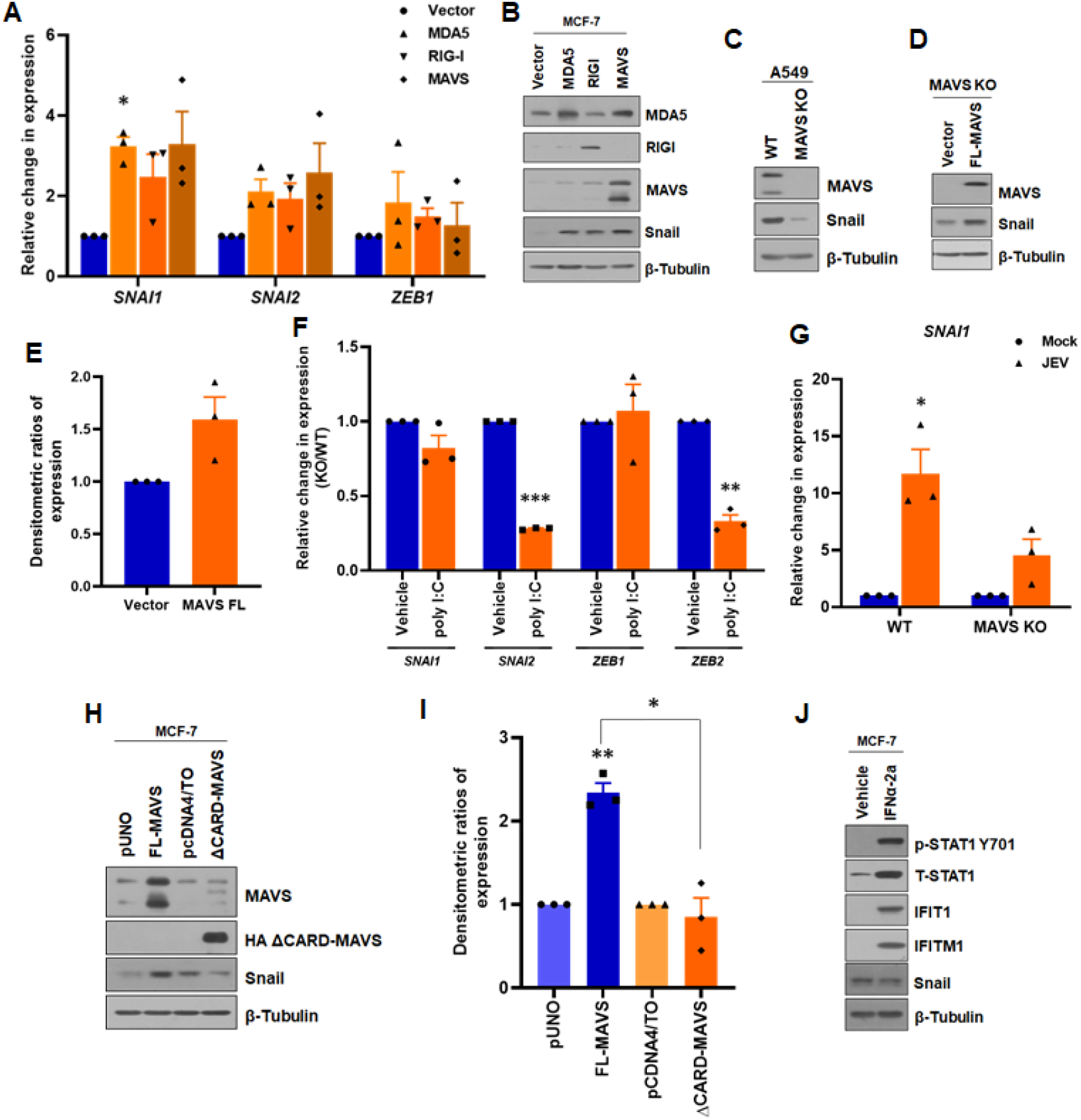
RLRs regulate EMT-TFs expression: (**A** & **B**) MDA5, RIG-I, and MAVS over-expression in MCF-7 cells. Real-time qRT-PCR of *SNAI1, SNAI2,* and *ZEB1* in over-expressed cells. (**A**) in comparison to vector-transfected cells. GAPDH was used as the normalization control. (**B**) Immunoblots for MDA5, RIG-I, MAVS, Snail, and β-Tubulin, confirming over-expression and EMT-TFs activation. (**C**) Snail expression in A549 MAVS KO was analyzed by immunoblotting. (**D** & **E**) Immunoblots of MAVS and Snail in MAVS KO A549 transfected with FL-MAVS construct. Densitometry. (**E**) for Snail expression normalized to β-Tubulin from three independent experiments. (**F**) Effect of poly (I:C) treatment on EMT-TFs, *SNAI1, SNAI2, ZEB1,* and *ZEB2* in MAVS KO A549, as compared to WT cells. (**G**) Effect on *SNAI1* in mock and JEV infected MAVS KO A549, in comparison with control cells. *GAPDH* was used as the normalization control. (**H**) Change in Snail expression upon ΔCARD-MAVS over-expression. ΔCARD-MAVS and full length (FL) were transfected into MCF-7 and checked for Snail by immunoblotting. (**I**) Densitometry of Snail expression normalized to β-Tubulin of the panel (**H**). (**J**) Immunoblots of IFNα-2a treated MCF-7 cells for the detection of EMT-TFs.

### Phosphorylated IRF3 promotes Snail transcription through ISREs

Thus far, our observations suggest that EMT-TFs are regulated by RLR-MAVS, but not by IFN. It is likely that this regulation might be downstream to MAVS, but upstream to IFN. Therefore, we focused our studies on IRF3, the key regulator in IFN signalling activated following RLR-MAVS activation. Ectopic expression of WT-IRF3 in MCF-7 cells failed to induce EMT-TFs. Nevertheless, phosphomimetic mutant S396E IRF3 (Figure 4A) efficiently activated both Snail and Slug expression reminiscent of viral infection (Figure 4B). A DNA binding mutant of IRF3 (DN-IRF3) also failed to induce the two molecules. S396E IRF3 activated *SNAI1* and *SNAI2,* but not *ZEB1* indicating that the regulation is possibly at the transcriptional level mediated by phosphorylated IRF3 (Figure 4C). In justification of these observations, only S396E IRF3 but not WT- and DN-IRF3 could strongly induce *IFNB1, IFIT1,* and *IFITM1* (Figure S4A). Consonant with this, we observed a significantly higher luciferase activity in Snail promoter-luciferase reporter vector (30) upon co-transfection with S396E-IRF3, but not with WT-IRF3 (Figure 4 D and E), confirming transcriptional regulation of *SNAI1* by p-IRF3.

**Figure 4:**
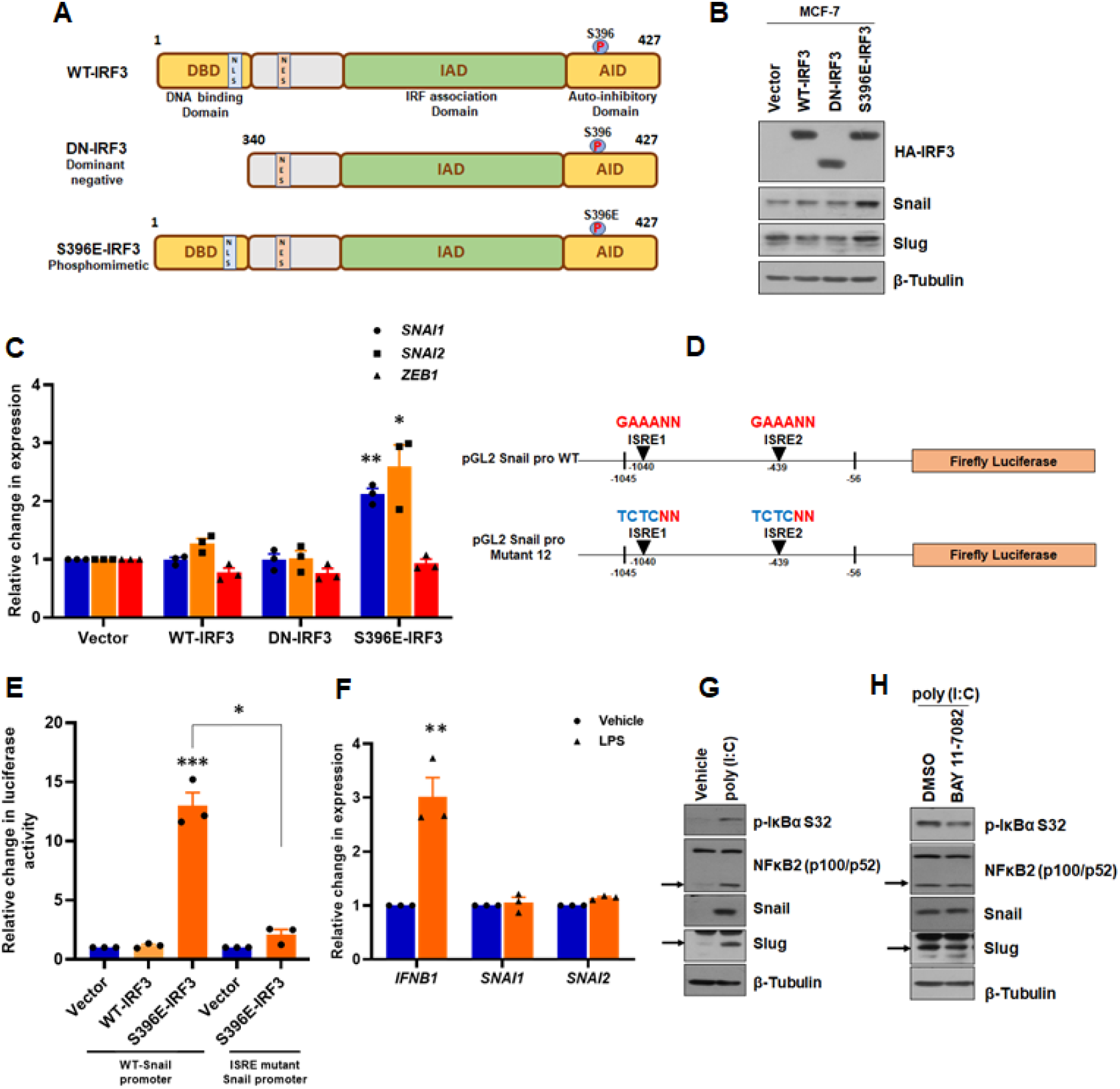
Phosphorylated IRF3 regulates *SNAI1* and *SNAI2* transcription: (**A**) Schematic representation of WT-IRF3 and its mutants DN and phosphomimetic S396E. (**B**) Immunoblot analysis of EMT-TFs in MCF-7 over-expressing the three IRF3 forms. (C) Quantitation of *SNAI1, SNAI2,* and *ZEB1* from panel (**A**) by real-time qRT-PCR, normalized to *GAPDH.* (**D**) Promoter analysis of Human Snail for putative ISRE in pGL2 Snail promoter construct. Two putative ISRE identified in the promoter region are indicated in the upper schematic while the lower demonstrates their mutant sequences pGL2 Snail ISRE mutant (GAAANN to TCTCNN) was created by site-directed mutagenesis at mentioned positions. The coordinates are with reference to TIS. (**E**) Dual-luciferase assay in MCF-7 cells co-transfected with WT- or S396E-IRF3 and WT or ISRE mutant Snail promoter-reporter constructs. pRL-CMV vector was used as the normalization control. Relative F/R ratios of luciferase activity are represented in the graph. (**F**) Real-time qRT-PCR of *IFNB1, SNAI1,* and *SNAI2* transcripts in LPS treated MCF-7 normalized to vehicle-treated cells. GAPDH was used as the normalization control. (**G**) NF-κB activation upon poly (I:C) treatment. MCF-7 cells were treated with 500 ng/mL poly (I:C) as mentioned in Figure 2. NF-κB pathway molecules were analyzed by immunoblotting, and β-Tubulin was used as the loading control. (**H**) Effect of NF-κB inhibitor, BAY 11-7082 on EMT-TFs. MCF-7 cells were treated with poly (I:C) for 22 hrs, followed by inhibition with 20 μM BAY11-7082 or DMSO for 2 hrs analyzed by immunoblotting for Snail and Slug.

IRF3 recognizes Type I IFN promoters through interactions with ISRE elements. To verify if IRF3 regulates SNAI1 through a similar mechanism, we performed sequence analysis of *SNAI1* promoter region and identified two potential consensus ISREs (GAAANN) at −1040 and −439 (Figure 4D). Double ISRE mutagenesis rendered the promoter-reporter non-responsive to phosphomimetic IRF3 and failed to activate luciferase (Figure 4E), suggesting that p-IRF3 activates *SNAI1* transcription by engaging with these elements. Based on these results, it is evident that viral RNA receptors RIG-I and MDA5 activate IRF3 that further promotes *SNAI1* transcription through ISREs in its promoter. Interestingly, our analysis of *SNAI2* promoter region identified four Type I (GAAANN) and two Type II (AANNNGAA) putative ISREs (Figure S4B), indicating its possible regulation through a mechanism similar as in *SNAI1.*

### NF-κB does not participate in *SNAI1* and *SNAI2* activation

Since NF-κB is a key transcriptional regulator in IFN production, we tested its requirement for EMT-TF transcription regulated by p-IRF3. TLR4 activation by LPS treatment caused phosphorylation of IκB (Figure S4C) and strong induction of *IFNB1* but failed to induce *SNAI1* and *SNAI2,* indicating that their activation by RLRs does not engage NF-κB (Figure 4F). Poly (I:C) activated NF-κB, as evident from phosphorylation of IκB and increased processing of precursor NF-κB2 (Figure 4G) in agreement with observations made earlier (31). However, NF-κB inhibitor BAY 11-7082 failed to suppress Snail and Slug levels in cells treated with poly (I:C) (Figure. 4H). These results demonstrate that NF-κB does not participate in *SNAI1* and *SNAI2* activation by viral RNA.

### Ectopic expression of EMT-TFs suppress RNA viral infections

Given that EMT, classically associated with tumor progression and embryogenesis, is activated by viruses using innate immune response pathway as demonstrated here, it is likely to have a consequence on the infection. To test this, we ectopically expressed Snail and Slug in A549 cells, followed by infection with JEV (Figure S5A). Expression of EMT-TFs imparted dramatic restriction to JEV infection evidenced by a significant drop in intracellular viral RNA and extracellular titers (Figure 5A) as well as expression of JEV envelope (Figure 5B). DENV infection was also substantially impacted by the EMT-TFs (Figure 5 C and D). The restriction on the virus was evident in MCF-7 cells as well (Figure S5 B-E). Importantly, EMT-TFs imparted only a limited suppression of viral entry (Figure S5F), indicating that the viral titer reduction is primarily associated with intracellular restriction.

**Figure 5:**
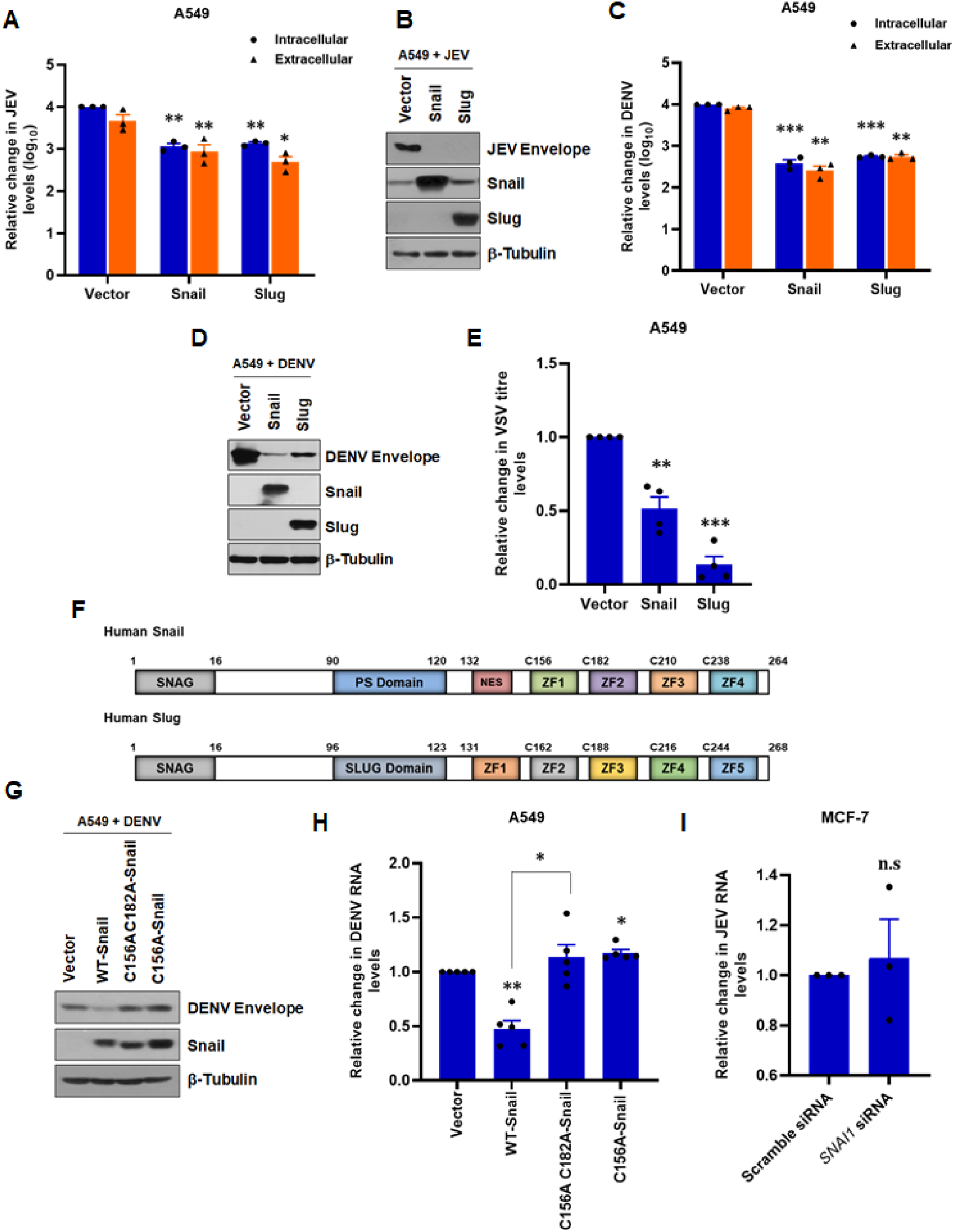
EMT-TFs suppress viral infections through its DNA binding activity: (**A**) A549 cells were transfected with Snail/Slug/empty vector, followed by JEV infection for 48 hrs. Supernatants and cells were harvested from infected cells and assessed for extracellular and intracellular titer by plaque assay and real-time qRT-PCR, respectively. (**B**) Immunoblot analysis of panel (**A**) samples tested for JEV envelope and confirmation of over-expression. (**C**) DENV infection on A549 cells transfected with EMT-TFs constructs, as mentioned in the experimental setup for (**A**). Extracellular titer measured by focus forming assay and intracellular titer by real-time qRT-PCR. (**D**) Confirmation of DENV infection and over-expression in samples from panel (**C**) by immunoblotting. (**E**) VSV infection of A549 cells transfected with Snail or Slug constructs. 18 hpi supernatant was collected and assayed for extracellular titer by plaque assay. (**F**) Schematic representation of DNA binding domains in Human Snail and Slug. (**G**) The importance of the DNA binding activity of Snail analyzed by immunoblotting of A549 cells expressing either WT- or double (C156A C182A) or single mutant (C156A)-Snail followed by DENV infection. (**H**) Real-time qRT-PCR of the panel (**G**) samples quantified for intracellular DENV RNA levels normalized to *GAPDH.* (**I**) MCF-7 cells were transfected with either 250 nM *SNAI1* siRNA or scramble using Lipofectamine 3000 for 24 hrs. Following transfection, cells were either mock or JEV infected at 0.1 MOI for 48 hrs. Cells were harvested and quantitated for intracellular JEV RNA levels by real-time qRT-PCR.

We then studied the universal efficacy of EMT-TFs as antiviral molecules by testing viral infection in diverse cell types upon their ectopic expression. Both Snail and Slug effectively conferred resistance against HCV in Huh7.5 cells (Figure S5G) and VSV in A549 cells (Figure 5E) marked by reduced viral RNA. These results comprehensively demonstrate that Snail and Slug promote antiviral state against RNA viruses. Next, we investigated if the transcription activity is critical for the antiviral activities displayed by Snail. Zinc fingers (ZF) are crucial to the DNA binding property of Snail (Figure 5F) and Slug, and mutations in ZF1 and ZF2 are known to suppress its transcriptional activity (32). As hypothesized, mutant Snail (C156A C182A) failed to impart any antiviral status to A549 cells against DENV infection, unlike its wild type form (Figure 5 G and H), indicating that Snail promotes its antiviral effects through transcriptional reprogramming. Thus, Snail and possibly Slug suppress viral infection through transcriptional reprogramming of genes involved in the antiviral program. To test if the depletion of one of the EMT-TFs supports viral infection, we knocked down Snail (Figure S5I) in MCF-7 cells using siRNA and subsequently infected with JEV. No significant change in JEV RNA levels (Figure 5I), could be observed upon the depletion of Snail. Since Snail and Slug exhibited antiviral activity individually when ectopically expressed, it is possible that absence of Snail could have been replaced by Slug. Hence, preferably a double knock out, Snail as well as Slug, is required.

### EMT-TFs activate innate immune response through RLRs

To find out the possible mechanism that contributes to the general virus restriction mediated by EMT-TFs, we analyzed their influence on the expression of various *ISGs*. Either of them activated *IFIT1* robustly and *IFIT3* modestly (Figure 6A). They also activated IFITMs *IFITM1, IFITM2,* and *IFITM3* (Figure 6B), indicating that IFN-mediated signaling activities were elevated. Interestingly, both *IFIH1* and *DDX58* were up-regulated by Snail and Slug (Figure 6C). Immunoblots confirmed these results and revealed increased phosphorylation of TBK1 and STAT1 in A549 cells expressing these EMT-TFs (Figure 6D). We reasoned that Snail and Slug could regulate *IFIH1* and *DDX58* as their promoter regions contained canonical E-boxes (Figure S6A). We verified the possibility of transcriptional regulation of *DDX58* by Snail and Slug by promoter-reporter assay. Their expression plasmids were co-transfected with a promoter-reporter construct of *DDX58* in A549 cells. However, no considerable change was observed in promoter activities, indicating that they do not regulate RIG-I transcriptionally (Figure S6B). We asked if similar results exist in published transcriptome studies since Snail and Slug activated innate antiviral signal pathways. Analysis of fourteen such datasets revealed that many ISGs are frequently up-regulated in EMT induced by diverse signals (Figure 6E). These results collectively indicate that EMT-TFs impart a strong innate antiviral response.

**Figure 6:**
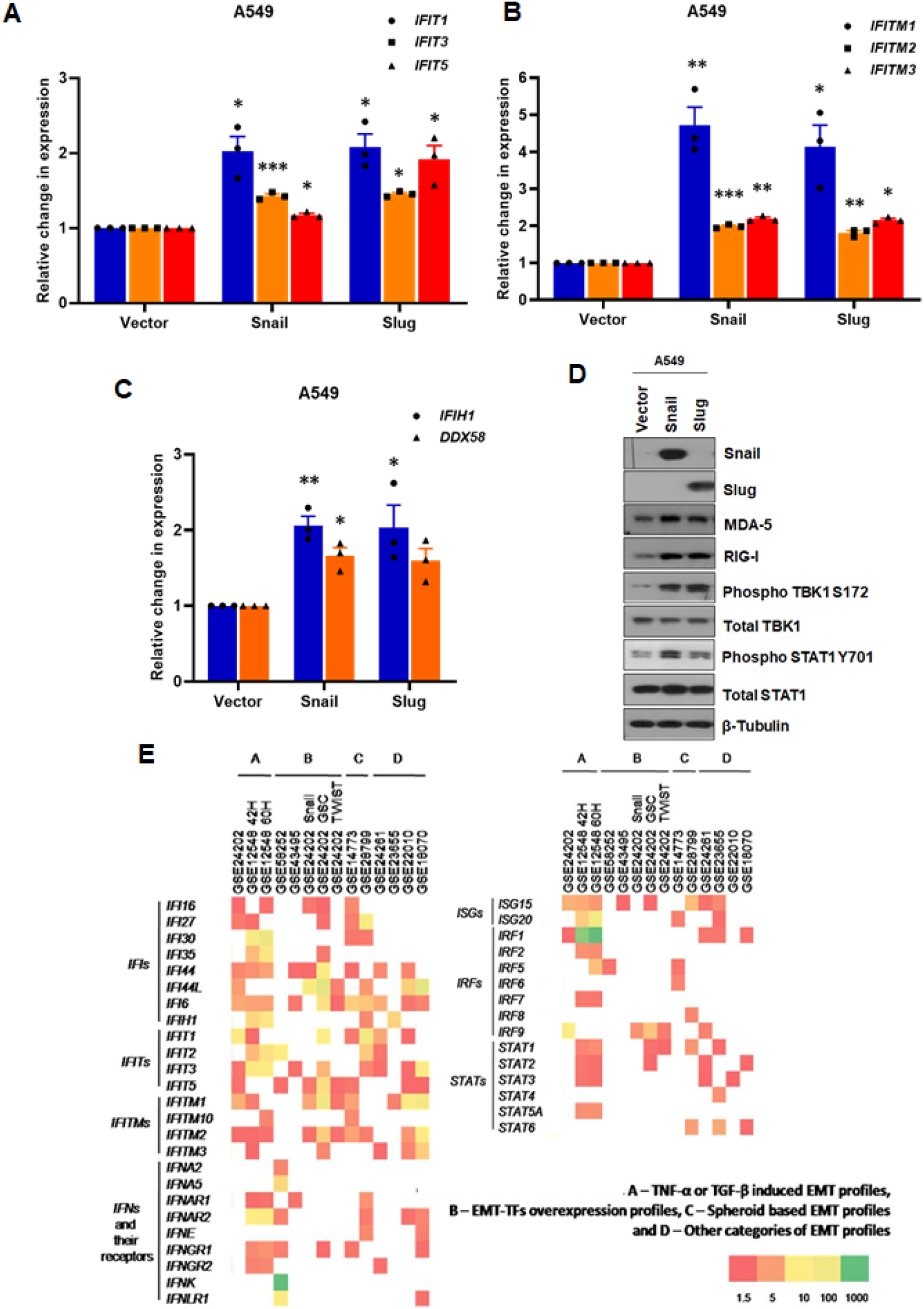
EMT-TFs activate cellular antiviral innate immune response: (**A-C**) A549 cells were transfected with Snail/Slug/vector using Lipofectamine 3000 and were harvested 72 hpt. Real-time qRT-PCR of IFITs, *IFIT1, IFIT3,* and *IFIT5* (**A**), IFITMs, *IFITM1, IFITM2,* and *IFITM3* (**B**), and RLRs, *IFIH1* and *DDX58* (**C**) in EMT-TFs over-expressed cells, normalized to *GAPDH.* (**D**) Cells from the above experiment were lysed, electrophoresed, and blotted for mentioned molecules. (**E**) Metadata analysis of fourteen EMT induced transcriptome profiles for cellular antiviral innate immune genes analyzed by GEO2R.

## DISCUSSION

EMT is most eloquently studied in cancer metastasis, embryo development, and wound healing. Despite the accumulated reports in several distinct conditions, the understanding of EMT during viral infections is minimal. Earlier reports focused on EMT induction during oncoviral infections, and these studies tried to establish a link between the process and oncogenesis promoted by these viruses. However, one paradox always stood out. If oncoviruses induce EMT early during infection in culture models, how would that have influenced cancer development that takes place several years after the infection? Further, mounting evidence suggests that EMT promotes metastasis in carcinoma (13). Detection of EMT or EMT-like process during infection of non-oncoviruses further indicated that EMT induced during viral infections could instead be a general host response that promotes a shared outcome. Our study initiated in this background revealed two essential aspects of EMT. First, it identified a novel mechanism of induction of two key EMT-TFs by a well-characterized pathway that is best known in coordinating the innate antiviral response. Second, it uncovered the consequence of the expression of EMT-TFs on viral infection by revealing their activation of antiviral responses. These findings have added a new dimension to our understanding of two physiological processes, EMT and antiviral response.

RIG-I and MDA5, two cytosolic RNA sensors, are crucial to alerting the cells against strange RNA signals in the cytoplasm. While RIG-I is known to have specificity for 5’-ppp carrying ssRNA or dsRNA, MDA5 primarily detects dsRNA. Since IVT generated HCV RNA with its uncapped 5’-ppp caused substantial induction of *SNAI1* and *SNAI2,* RIG-I is an effective inducer in this process. On the other hand, MDA5 senses double-stranded replicative forms common during the phase of active replication, and genomes of dsRNA viruses. Thus, the evolutionary process has ensured redundancy in this process. Interestingly, a mechanism that has evolved to activate Type I IFN is also shared to activate EMT-TFs, which in turn, perpetuate the mechanism itself. Equally impressive is the specificity towards *SNAI1* and *SNAI2,* leaving out *Twist* and *Zeb1* from this ambit.

It is interesting to note that the depletion of Snail did not have any impact on the viral titer which was expected under normal circumstances considering its antiviral effect. In agreement, the ectopic expression of the DNA binding mutant of Snail also had little effect on the viral titer. The most plausible explanation for these observations is the functional redundancy of EMT-TFs. It is very likely that Slug and possibly the other members of EMT-TFs are compensating for the loss of Snail, effectively suppressing the viral replication. Thus, demonstrating more complete roles of these transcription factors in viral infection would necessitate a multiple knockout system.

Our model proposes that EMT-TFs form a loop that helps sustain the antiviral response through RLRs (Figure 7). While we do not yet completely understand this process’s mechanistic details, we demonstrate that EMT-TFs can modulate the levels of RIG-I and MDA5. Since it does not appear to be a transcriptional regulation, other possibilities, such as stabilizing the proteins through post-transcriptional means exist. Ubiquitination is a well-known mode of activation of both RIG-I and MDA5 (33, 34) and appears to be a strong possibility. Importantly, the upstream circuit of regulating Snail and Slug expression as well as that downstream to it regulating RLRs are exclusive to the infected cells as extraneous IFN had very little effect on the EMT-TFs. It is interesting to note that the mere expression of these molecules in naïve cells was enough to induce IFN as well as EMT-TF transcriptionally.

**Figure 7:**
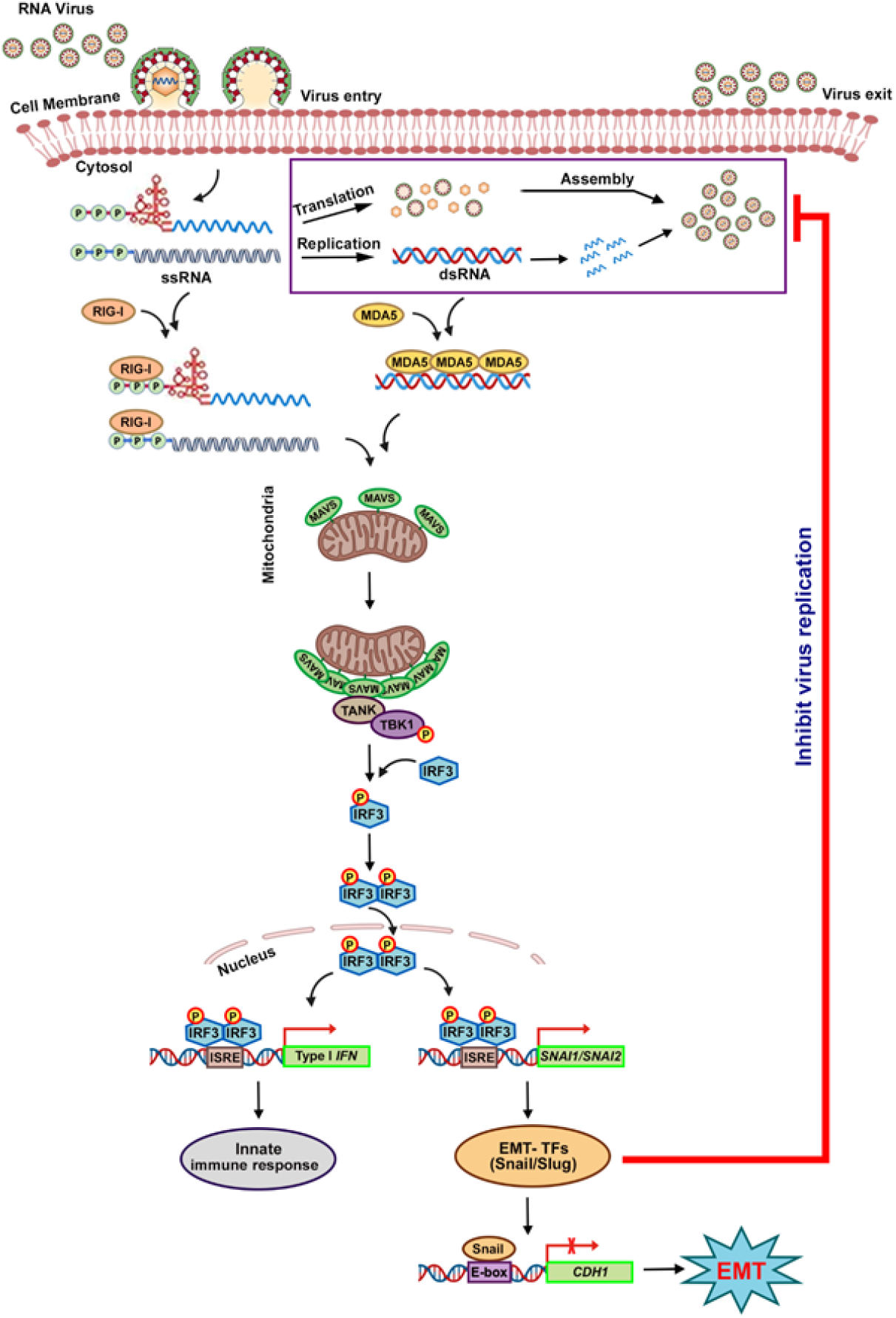
Model illustrating a novel mechanism for activation of EMT-TFs during RNA viral infections: After the virus entry into the cell either through endocytosis or fusion, the viral genome is sensed by RLRs, RIG-I, and MDA5, which transmits signals to MAVS localized on mitochondria, leading to oligomerization of MAVS. This recruits TANK and TBK1 (TANK binding kinase 1) onto oligomerized MAVS, resulting in activation of TBK1 by TANK through phosphorylation at S172. Phosphorylated TBK1 activates IRF3 through a series of phosphorylations at the C-terminus of IRF3, resulting in dimerization and subsequent nuclear translocation of IRF3, leading to transcriptional activation of *SNAI1* and possibly *SNAI2* as in the case of Type I IFN promoter activation. Snail and Slug, on the one hand, elevates ISGs and, on the other, represses E-cadherin, resulting in EMT. Elevated antiviral genes by Snail and Slug restrict viral replication, thereby controlling the spread of infection.

IRF3 is a common node for both RNA and DNA viruses. A major mechanism of antiviral response to DNA viruses is coordinated by the cGAS/STING pathway that senses DNA in the cytoplasm and activates IRF3 (7, 35), further leading to IFN production. Thus, it is very likely that one of the significant mechanisms of EMT activation during DNA viral infection involves this pathway, with IRF3 being the common molecule that facilitates the induction of Snail and Slug during both RNA and DNA viral infections. We tested this hypothesis using EBV, a γ-herpesvirus with dsDNA as genome that displays a clear tropism for both epithelial and B cells. Interestingly, the EMT-TFs displayed a strong antiviral effect against EBV, as measured by decreased *LMP1* transcripts (Figure S5H). However, whether sensing of DNA by cGAS/STING activates EMT-TFs is an open question. This model suggests that viral genomes, whether RNA or DNA, are potent inducers of EMT-TFs that play important roles in sustaining antiviral response. Therefore, the constant presence of viral genomes in persistent viral infections could result in prolonged expression of EMT-TFs that might, in turn, collaborate with other factors in promoting cancers in such cases. However, this does not rule out the possibility of other well-established mechanisms of EMT-TF induction in parallel. This includes viral proteins that could separately influence pathways such as TGF-β and Wnt/β-Catenin. Thus, these complex networks of pathways influencing each other in dynamic ways would decide the outcome of infection.

Type I IFNs are the primary target genes of IRF3 known so far. The discovery that the latter can regulate *SNAI1* (and most likely *SNAI2*) transcriptionally opens up the possibility of a much larger role of IRF3 in various other biological events. Recent studies linking the potential antagonistic effects of RIG-I in cancer progression (36, 37) should be analyzed in the context of our studies to understand the broader implications of EMT-TFs expressed under these conditions. Over-expression of IRF3 and its mutants are reported in several cancers (38–40). Increased activation of TBK1 (TANK binding kinase 1) and IKKε has been reported in various cancers (38). On a different note, DNA released from tumor cells or dead cells can enter neighboring antigen-presenting cells through unknown mechanisms that can activate STING in them (41). Cancer cells are well known to modulate their immune molecules in their quest to avoid immune surveillance, and the contribution of EMT to this cause is well studied (42). In this context, a leading role played by IRF3 in determining the outcome of the complex signaling networks is conceivable. Certainly, concurrent regulation and expression of anti-cancerous IFN and pro-cancerous EMT-TFs looks intriguing and demands more detailed studies. The activation of RLR signaling by Snail and Slug further suggests that they might assist IRF3 in suppressing cancer progression. Further studies into the factors that determine the pro-growth or anti-growth effect of these two molecules are necessary. Nevertheless, a larger role of IRF3 in the overall progression and survival of cancer cells is quite likely, and our studies provide important clues for the long standing question in the field of cancer biology.

Apart from the two mentioned contributions, our study raises a few critical questions. Our studies have primarily dealt with EMT-TFs in epithelial cells. However, non-epithelial cells also express EMT-TFs, and the RLR pathway is fairly conserved across cells, including specialist cells, and it is likely that the mechanism that we identified is more general and not restricted to epithelial cells. Another point of future interest is the probably elevated antiviral state in circulating tumor cells (CTCs) in developing embryos and stem cells. CTCs regulate the expression of their cell surface receptors in order to evade immune surveillance. Elevated antiviral state imparted by EMT-TFs could further protect them from possible viral infections, thereby increasing their probability of survival in circulation. Interestingly, stem cells are known to be refractory to various viral infections, and the contribution of EMT-TFs to this status needs special attention. EMT could be a mechanism that could help the cells alter the tropism of viruses. Here, EMT mediated changes in the expression of various host factors could enforce stringent survival conditions on the virus and could be a ploy by the cells to limit the spread of the virus. This point is particularly strengthened by the absence of HCV in liver tissues with higher tumorigenic indices (43). Compromised infection of transited mesenchymal cells by Measles virus also points in this direction (44). A partial drop in virus entry following the expression of EMT-TFs in our studies also suggests that EMT might be accompanied by altered receptor expression rendering a more impermissivity to the cells.

Based on the functional outcome in viral infection, which is distinct from the other established EMTs, we propose virus-induced EMT as the fourth paradigm of EMT that attributes a fresh dimension to EMT. While the RLR-IRF3 axis may activate EMT in the other three contexts, its activation is associated with viral infections where it provides a unique functional outcome. Future studies would provide details of the interplay between the innate immune signaling and other biological events such as cancer metastasis.

## AUTHORS CONTRIBUTION

D.V and K.H.H conceived and designed the project. D.V performed, analyzed, and interpreted the results of experiments using JEV, DENV and HCV. D.G performed viral entry and IFN treatment experiments and interpreted the results. Initial experiments on DENV and JEV were performed in the laboratory of M.K. M.B and A.B designed and executed mice experiments. D.V and K.H.H analyzed the results of mice experiments. G.K and M.V.V performed and analyzed EBV related experiments. D.G, A.M, and D.N executed and interpreted VSV related experiments. D.V assisted K.H.H to write the manuscript.

## ACKNOWLEDGEMENTS

Special thanks to Mohan Singh Moodu and Amit Kumar for maintaining the logistics needed for the entire work. We are thankful to Sankar Bhattacharyya for his assistance with establishing DENV cultures. We thank the contributions made by Vishal Sah, Hitha G Nair, Poonam Manral and Sriram Kumar in standardization-related experiments. Mohan Singh Moodu, Farsana S.M. Swetha Jeevan, and Karthika S Nair are acknowledged for cloning experiments. pUNO vector, pUNO RIG-I, pUNO MDA5, and pUNO IPS-1 (MAVS) were a kind gift from Dr. C T Ranjit Kumar, THSTI.

## MATERIALS AND METHODS

### Antibodies, reagents, plasmids, cloning, and cell lines

All primary antibodies were purchased from Cell Signaling Technologies, except anti-flaviviral envelope 4G2 (Merck Millipore), β-Tubulin, and GAPDH (Thermo Fischer), and HCV Core (Abcam). Rabbit polyclonal JEV NS1 antibody was from Dr Manjula Kalia, RCB, Faridabad. All HRP-conjugated secondary antibodies were procured from Jackson Immunoresearch. Poly (I:C), BAY 11-7082, PEI, and Human IFNα-2a were purchased from Sigma Aldrich. *SNAI1* and Scramble siRNA were purchased from Dharmacon, Thermo Scientific.

Mammalian expression vector, pcDNA4/TO, was procured from Invitrogen. All over-expression constructs were generated using pcDNA4/TO. HA-IRF3 was amplified using primers and MCF-7 cDNA as template to generate pcDNA4/TO IRF3. Similarly, the DN-mutant was amplified with specific primers and cloned. Phosphomimetic IRF3 (S396E) was generated using primers carrying corresponding mutations by overlapping PCR. HA-ΔCARD-MAVS was amplified using pUNO MAVS as template and cloned into pcDNA4/TO. pcDNA4/TO Snail and pcDNA4/TO Slug were generated by amplifying their respective cDNA clones procured from Thermo Fischer. Zinc finger single (C156A) and double (C156A C182A) mutants of Snail were generated using pcDNA4/TO Snail as template by overlapping PCR. Snail_pGL2 was a gift from Paul Wade (Addgene plasmid # 31694; http://n2t.net/addgene:31694; RRID: Addgene_31694). Snail promoter-luciferase reporter construct was used as template to generate pGL2 Snail ISRE mutant promoter construct. pGL2 DDX58 (RIG-I) promoter-reporter was generated using primers and MCF-7 genomic DNA as template. All constructs generated in the lab were sequenced to confirm their integrity. All primers used in the study for generating constructs are listed in Table 1. Plasmids were prepared from bacterial cells using MN Nucleospin Plasmid DNA kit.

**Table 1:**
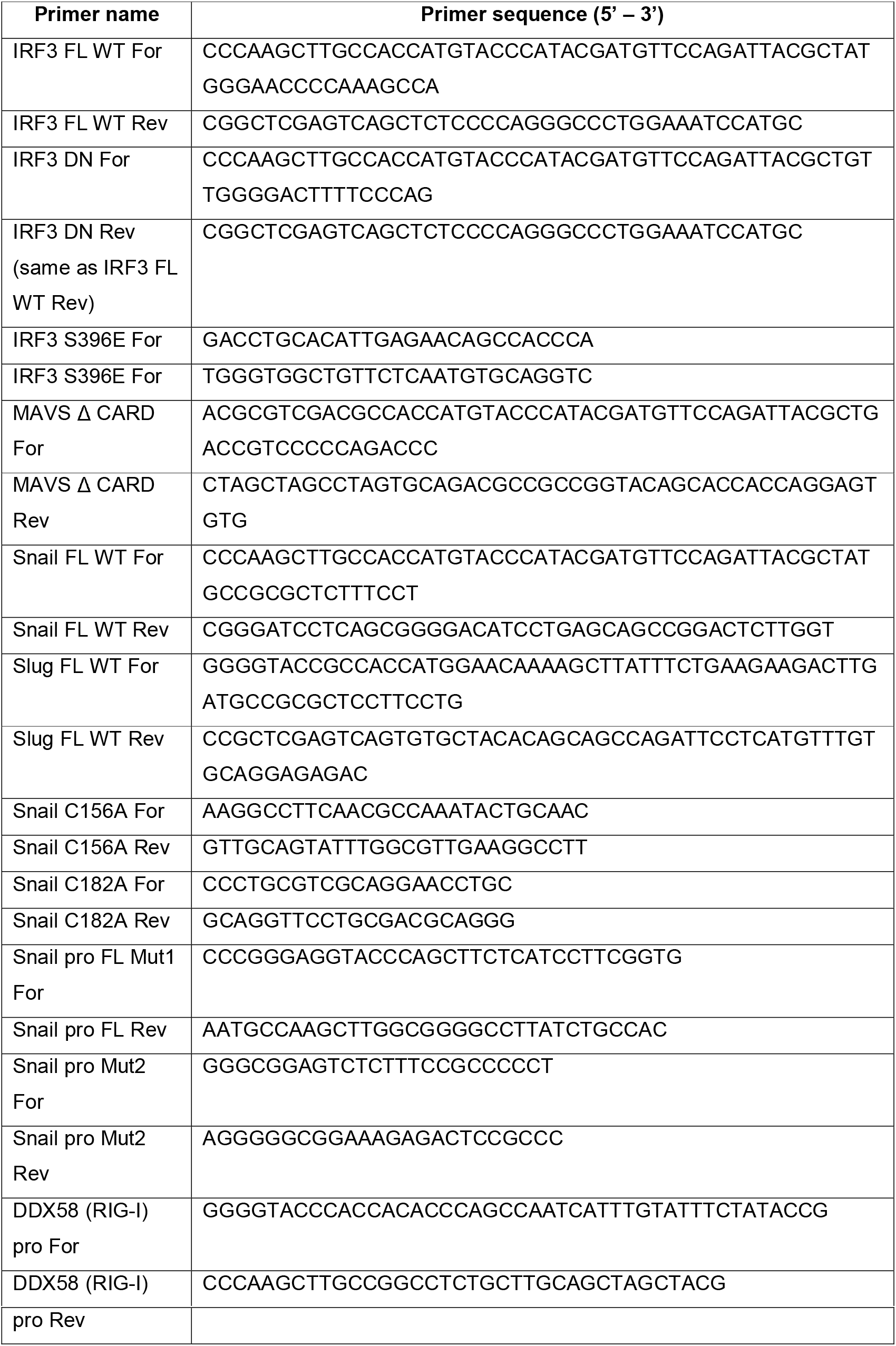
List of primers used for generating constructs.

MCF-7, A549, and HEK293 were received from ATCC. Vero and PS cells were received from Dr. Manjula Kalia, RCB, Faridabad. All cells mentioned above were cultured in DMEM supplemented with 10% FBS, 1X Penicillin-streptomycin, and grown at 37 °C with 5% CO_2_. Huh7 and Huh7.5 were gifted by Dr. Ralf Bartenschlager and Dr. Charlie Rice, respectively. They were also cultured in the above media, along with 1X NEAA supplement. A549 Dual™ WT and A549 Dual™ MAVS KO (MAVS KO) cells were procured from Invivogen and cultured according to manufacturer’s protocol. These cells were used in experiments pertaining to the comparison between MAVS KO and its control. All cell lines were tested negative for mycoplasma by PCR using specific primers (Table 2).

**Table 2:**
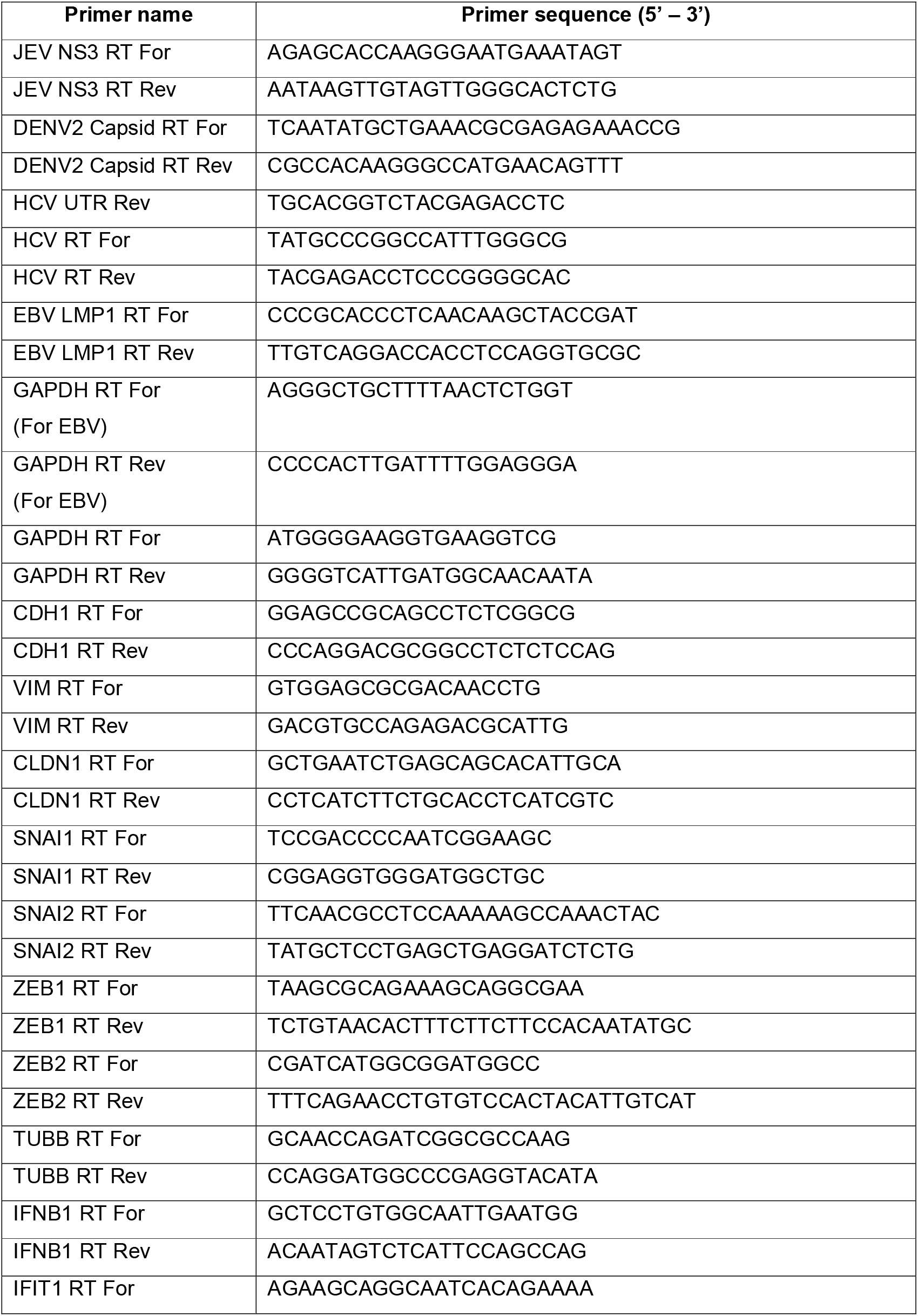
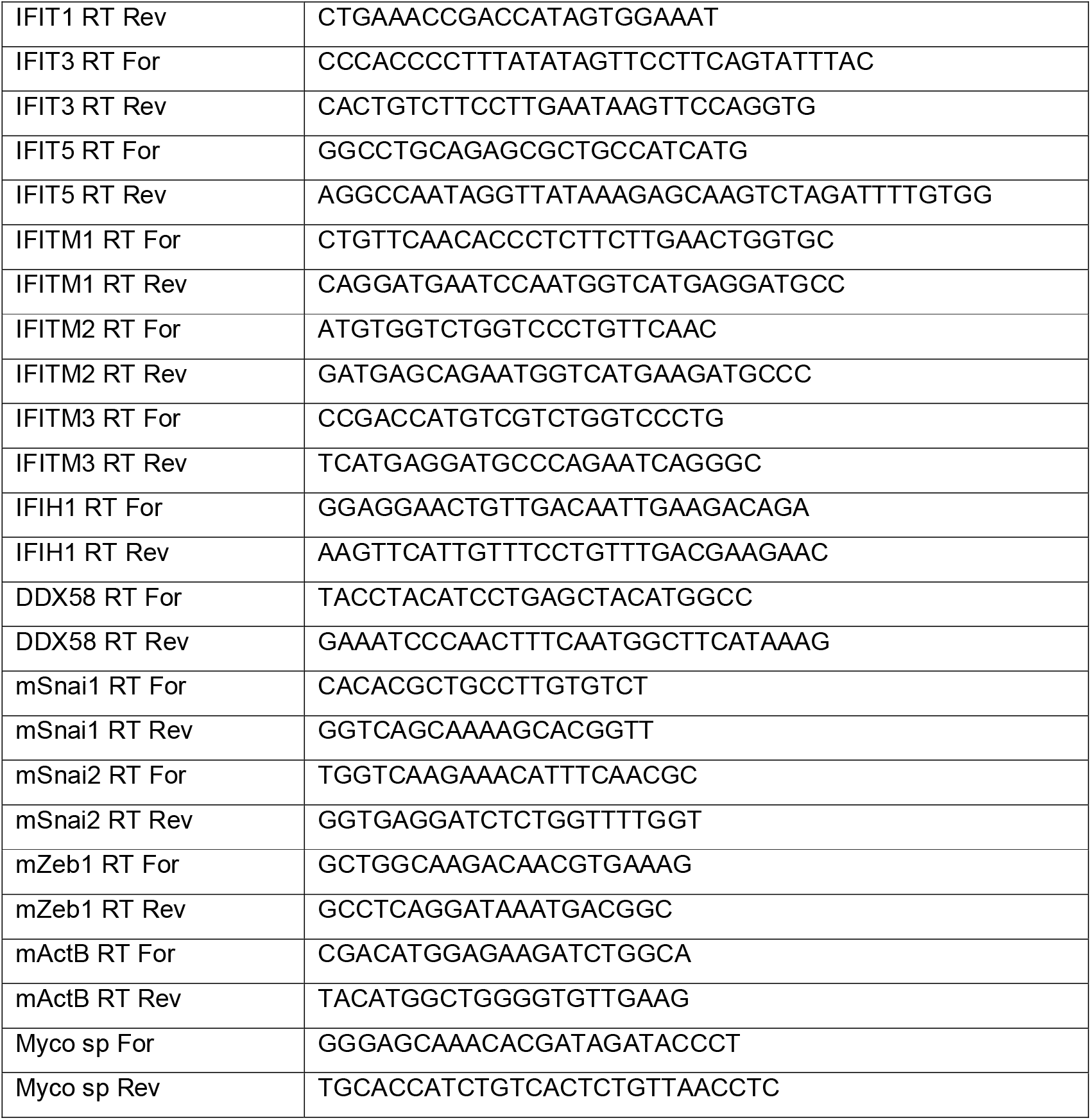
List of primers used for real-time qRT-PCR

### Propagation and purification of viruses

All infection-related experiments were carried out in Bio Safety Laboratory (BSL) level – 2 at CSIR-CCMB, Hyderabad. Infectious JEV isolate (Vellore strain P20778, genotype 3, GenBank accession No. AF080251) was a kind gift from Dr. Manjula Kalia, RCB, Faridabad (45). JEV was propagated in Huh7 cells and incubated for 36-48 hrs. On observation of the cytopathic effect, the culture supernatants of JEV infected cells were collected and spun at 3500 rpm for 10 min at 4 °C. Cell-free supernatant was passed through 0.2 μm filter, aliquoted, and stored at −80 °C. Plaque assay was performed to determine the viral titer.

Dengue virus serotype 2 (DENV2) was a kind gift from Dr. Sankar Bhattacharyya, THSTI, Faridabad (46). Like JEV, DENV was also propagated and stored at −80 °C. Foci forming assay was performed to determine DENV titer. Hepatitis C Virus 2a was propagated in human hepatoma cell line, Huh7.5 (47, 48). HCV titer was calculated by real-time qRT-PCR using the absolute quantification method. EBV was prepared from B95-8 cells by inducing the lytic cycle in the BSL-2 laboratory of Dr. Mohanan Valiya Veettil (49). Briefly, B95-8 cells were treated with 20 ng/mL of TPA and incubated for 5 days at 37 °C. The supernatant was filtered and ultracentrifuged at 70000 rpm for 2 hrs at 4 °C, and the viral pellet was dissolved in PBS, and the virus titer was determined (50).

Vesicular stomatitis virus (VSV) (Indiana strain) was a kind gift from Dr. Debasis Nayak, IIT Indore. VSV was propagated in Vero cells at 0.01 MOI. When 50% cytopathic effect (CPE) was observed, the culture supernatant was collected and centrifuged at 1400 × g for 10 min at 4 °C. Debris free supernatant was filtered through a 0.2 μm membrane filter. The clarified supernatant was then aliquoted and stored at −80 °C. Plaque assay was performed to determine the VSV titer.

### Infection and quantification of viruses

For JEV infection, cells were seeded to reach 60% confluency. Infection was carried out using DMEM without antibiotics. For JEV infection, cells were infected with JEV at 0.1 MOI for 2 hrs in DMEM. Simultaneously, the mock infection was also set. 2 hpi, the inoculum was removed, and cells were washed twice with PBS and replenished with cDMEM containing 10% FBS, 1X Penicillin-Streptomycin, and incubated at 37 °C for 48 hrs. Plaque assay was carried out to determine the JEV titer (51). Briefly, PS cells were seeded in a 6-well plate to form monolayer. Serial dilution of the virus was performed in DMEM. The monolayer was inoculated with serially diluted virus inoculum and incubated at 37 °C for 1 hr with gentle shaking. After incubation, the inoculum was removed and washed once with PBS. The infected monolayer was overlaid with overlay media (2X cDMEM and 2% Low melting agarose (LMA) in 1:1 ratio). The plates were incubated for 3-5 days until plaques were visible. Cells were fixed with 10% formaldehyde and stained with Crystal violet. Plaques were counted from each dilution, and the titer was calculated and represented as pfu/mL.

Similarly, for DENV infection, 0.5 MOI was used, and cells were incubated with inoculum for 4 hrs, and the protocol mentioned above was followed. Mock and DENV infected cells were incubated at 37 °C for 72 hrs. Cells and supernatant were collected post-infection. Foci forming assay was carried out to determine DENV titer (46). Vero cells were seeded in a 12-well plate to form monolayer. The serially diluted virus was inoculated onto monolayers and incubated at 37 °C for 4 hrs with gentle shaking. The inoculum was removed, and overlay media (2X cDMEM and 2% Carboxymethylcellulose (CMC) in ratio 1:1) was added to cover the monolayer, and the cells were incubated at 37 °C for 5-6 days. Overlay media was removed, and cells were fixed with 4% Formaldehyde. Cells were washed with PBS to remove Formaldehyde traces and permeabilized with PBS containing 0.1% Triton-X 100. Cells were then blocked with 2% BSA prepared in PBS at room temperature for 1 hr. Cells were then incubated with mouse anti-flaviviral specific envelope antibody (4G2) (1:2000) overnight at 4 °C. Following this, HRP-conjugated anti-mouse antibody (1:5000) was added to cells and incubated at room temperature for 2 hrs. After three washes with PBS, the foci were developed using TMB substrate. Foci were developed and visualized under white light illuminator. Foci were counted, and the DENV titer was calculated as represented as ffu/mL.

For determining HCV titer in the supernatant, RNA was prepared by the Trizol method from cell-free viral supernatant. cDNA was prepared by reverse transcription using Primescript Reverse transcriptase (Takara) and HCV specific RT primer (Table 2). 5’UTR of HCV 1b was used as the standard to determine the HCV copies present in the supernatant. Quantitative real-time PCR was set for prepared cDNA and serially diluted standard (copy number was calculated and diluted from 10^8^ to 10^3^) using SYBR Green mastermix (Takara) and HCV specific PCR primers in LightCycler 480 instrument (Roche). By absolute quantification method, a standard curve was prepared for standards, and copy number of the unknown sample was calculated and represented as copies/mL (52).

For VSV infection, cells were seeded and infected with VSV at 0.1 MOI in PBS with MgCl_2_ and CaCl_2_ for 1 hr with shaking every 10 minutes. Virus inoculum was removed, and cDMEM was added. Cells were incubated at 37 °C for 18 hrs. All mock and virus-infected cells and their supernatants were harvested at specified time points and collected for RNA or protein work. VSV titer was determined by plaque assay with a slight modification in the protocol used for JEV. Vero cells were seeded in a 6-well plate to form confluent monolayer. Serially diluted virus inoculum in PBS with MgCl_2_ and CaCl_2_ for 1 hr with shaking at every 10 minutes. The inoculum was removed and washed with PBS. Cells were overlaid with 0.8% overlay media (2X cDMEM and 1.6% low melting agarose) and incubated at 37 °C for three days until clear plaques were visible. Cells were fixed with 10% Formaldehyde, followed by the removal of overlay media. Fixed cells were then washed with PBS and stained with Crystal violet stain (0.1% Crystal violet, 0.65 g Na_2_HPO_4_, 0.4 g NaH_2_PO_4_, 90 ml H_2_O, 10 ml 37% Formaldehyde). Once plaques are visible, the stain was removed, and plates were washed with running water. Plaques were counted, and titer was represented as pfu/ml.

### JEV infection in mice

All *in vivo* experiments were performed on ten-day-old BALB/c mice who were always housed with their mothers under pathogen-free and climate-controlled conditions with 12-hrs light/dark cycle at a constant temperature of 25 ± 2 °C and relative humidity 60 ± 10%. For experiments, pups of either sex were randomly divided into two groups: Sham-treated and JEV-infected. Mice belonging to the JEV-infected group were injected with 3 x 10^5^ PFU of GP-78 strain intraperitoneally, while an equal amount of sterile PBS was injected into the sham-treated group (53). Infected pups, along with sham-treated animals, were euthanized on day seven post-infection after the encephalitis symptoms appeared (54, 55). Brain and liver were excised after transcardial perfusion of anesthetized animals with ice-cold 1X PBS, and thus obtained tissue samples were used for RNA work. All experiments conducted were approved by the Animal Ethics Committee of National Brain Research Centre (Approval no - NBRC/IAEC/2017/130) and were in accordance with the guidelines of the Committee for the Purpose of Control and Supervision of Experiments on Animals (CPCSEA), Ministry of Environment and Forestry, Government of India.

### Transfections, treatments, and infection

For poly (I:C) transfections, cells were seeded to reach 80% confluency. Transfection mix containing OptiMEM-Lipofectamine 3000-poly (I:C) was prepared according to the manufacturer’s protocol and added to cells and incubated for 6-8 hrs. After incubation, the transfection mix was removed, and cDMEM was added and further incubated for 16-18 hrs. 24 hpt, mock (PBS), and poly (I:C) transfected cells were harvested and collected for RNA or protein work.

For the poly (I:C) and inhibition experiment, the above protocol was followed. After 22 hrs of poly (I:C) treatment, spent media was removed. cDMEM supplemented with DMSO or BAY 11-7082 (20μM) was added to poly (I:C) transfected cells continued for 2 hrs accounting for a total of 24 hrs of incubation.

For LPS treatment, cells were seeded to reach 80% confluency. Spent media was removed, and fresh cDMEM containing either PBS (vehicle) or LPS (1 μg/mL) was added and incubated for 6 hrs. Cells were harvested and used for RNA or protein studies.

For IFNα treatment, cells were seeded to reach 70% confluency. Spent media was removed, and fresh cDMEM containing either PBS (vehicle) or 50 ng/mL of IFNα was added and incubated for 24 hrs. Cells were harvested and used for RNA or protein work.

For EMT-TFs over-expression and infection studies, Cells were seeded to reach 60% confluency. Snail and Slug over-expression vectors, along with empty vector, were transfected using Lipofectamine 3000 according to the manufacturer’s protocol. 8 hrs post-transfection, the transfection mix was removed, and cDMEM was added. 24 hpt, cells were infected with viruses at mentioned MOI. JEV at 0.1 MOI, DENV at 0.5 MOI, HCV at 0.5 MOI, and VSV at 0.1 MOI. 48 hpi, mock, and virus-infected cells were harvested, except for HCV at 72 hpi. Viral supernatants were collected to determine the extracellular titer as mentioned above, while intracellular titer was quantified by real-time qRT-PCR using virus-specific primers (Table 2).

For EBV, an equal number of cells were plated the day before transfection. Cells were transfected with over-expression constructs using 1 μg/μL PEI (Sigma) and incubated for 5 hrs at 37 °C with subsequent media change. Transfected cells were cultured for 48 hrs and then infected with EBV for 2 hrs. 48h post EBV infection, cells were harvested for RNA isolation.

### Immunoblotting

For protein work, cells harvested were washed with ice-cold PBS supplemented with 1mM PMSF. Cells were lysed using a mild NP-40 lysis buffer and incubated on ice for 10 min with gentle mixing. Lysed samples were centrifuged at 13000 rpm for 15 min at 4 °C. The clarified supernatant was collected and used as a crude protein lysate. All lysates were quantified by BCA method. Equal quantities of lysates were loaded on SDS-PAGE, electrophoresed, and transferred onto activated PVDF membrane by wet transfer method. The membrane was blocked and probed for respective molecules with primary and HRP conjugated secondary antibodies. Blots were developed by classical method autoradiography. For DENV and JEV envelope protein, the lysates were mixed with sample loading dye without β-Mercaptoethanol and loaded onto SDS PAGE.

### cDNA synthesis and real-time quantitative RT-PCR

For all RNA work, cells were harvested were washed with ice-cold PBS. Total RNA was isolated using MN Nucleospin RNA kit (Takara). Equal quantities of RNA were reverse transcribed using Primescript Reverse transcriptase and random hexamers (Takara). 50-100 ng of cDNA was used for quantification using SYBR Green mastermix (Takara) in Lightcycler 480 instrument (Roche). All virus and host-specific transcripts were normalized to GAPDH. By relative quantification method, ΔΔ Cp (crossing point) was calculated, and fold change (2^(ΔΔCp)^) in comparison to vector-transfected and infected control cells is represented in the graphs.

For EBV quantification, total RNA from Snail and Slug transfected, and EBV infected A549 cells was extracted using TRI Reagent (Sigma) and treated with DNase (Promega). Equal quantities of total RNA were used for cDNA synthesis using High Capacity cDNA reverse transcription kit (Applied Biosystems). Real-time quantitative RT-PCR was performed using EBV *LMP1* gene-specific primers and *GAPDH,* Power SYBR green mastermix (Applied Biosystems) in Applied Biosystems 7300 Real-Time PCR instrument. Fold change was calculated as mentioned above. The list of primers used in qRT-PCR is given in Table 2.

### Luciferase assay

For luciferase reporter assays, cells were co-transfected with either IRF3 WT or IRF3 S396E and Snail promoter WT/ISRE mutant reporter construct, along with pRL-CMV vector as normalization control using Lipofectamine 3000 (Invitrogen). 30 hpt, cells were harvested and proceeded with Dual luciferase assay (Promega) according to the manufacturer’s protocol in a luminometer. Firefly and Renilla luminescence readings were noted. All Firefly readings were normalized with respective Renilla reading. The obtained (F/R) ratio was normalized to empty vector-transfected F/R ratio, and relative change in the F/R ratio was calculated.

For DDX58 promoter assay, Snail/Slug over-expression constructs were co-transfected with pGL2 DDX58 Firefly luciferase reporter construct and pRL-CMV vector as mentioned above. The relative F/R ratio calculated was used to make a graphical representation.

### Metadata analysis of transcriptome profiles

For metadata analysis of transcriptome profiles, fourteen different EMT induced transcriptome profiles were retrieved from GEO database. TNF or TGF-β induced profiles (GSE24202, GSE12548 42 hrs, and GSE12548 60 hrs), EMT-TFs over-expression profiles (GSE58252, GSE43495, GSE24202_Snail, GSE24202_GSC and, GSE24202_TWIST), spheroid based EMT profiles (GSE14773 and GSE28799), and other all category EMT profiles (GSE24261, GSE23655, GSE22010 and, GSE18070) were used for this metadata analysis. GEO2R was used to identify genes involved in cellular innate antiviral immune response with at least 1.5-fold and p-value <0.5. Such a list of genes was prepared from all fourteen profiles, and heatmap was generated.

### Graphs and statistical analysis

Statistical significance was calculated by the paired-end, two-tailed Student’s t-test method. All experiments were conducted in a minimum of three independent rounds, and averaged values are represented as scatter plots with bar graphs (depicting individual values of independent experiments). Error bars are representations of the mean ± SEM. All graphs were prepared using GraphPad Prism version 8.0.2. Statistical significance is represented as *, **, and *** for p<0.05, p<0.01 and p<0.005 respectively.

## SUPPLEMENTARY FIGURES

**Figure S1:**
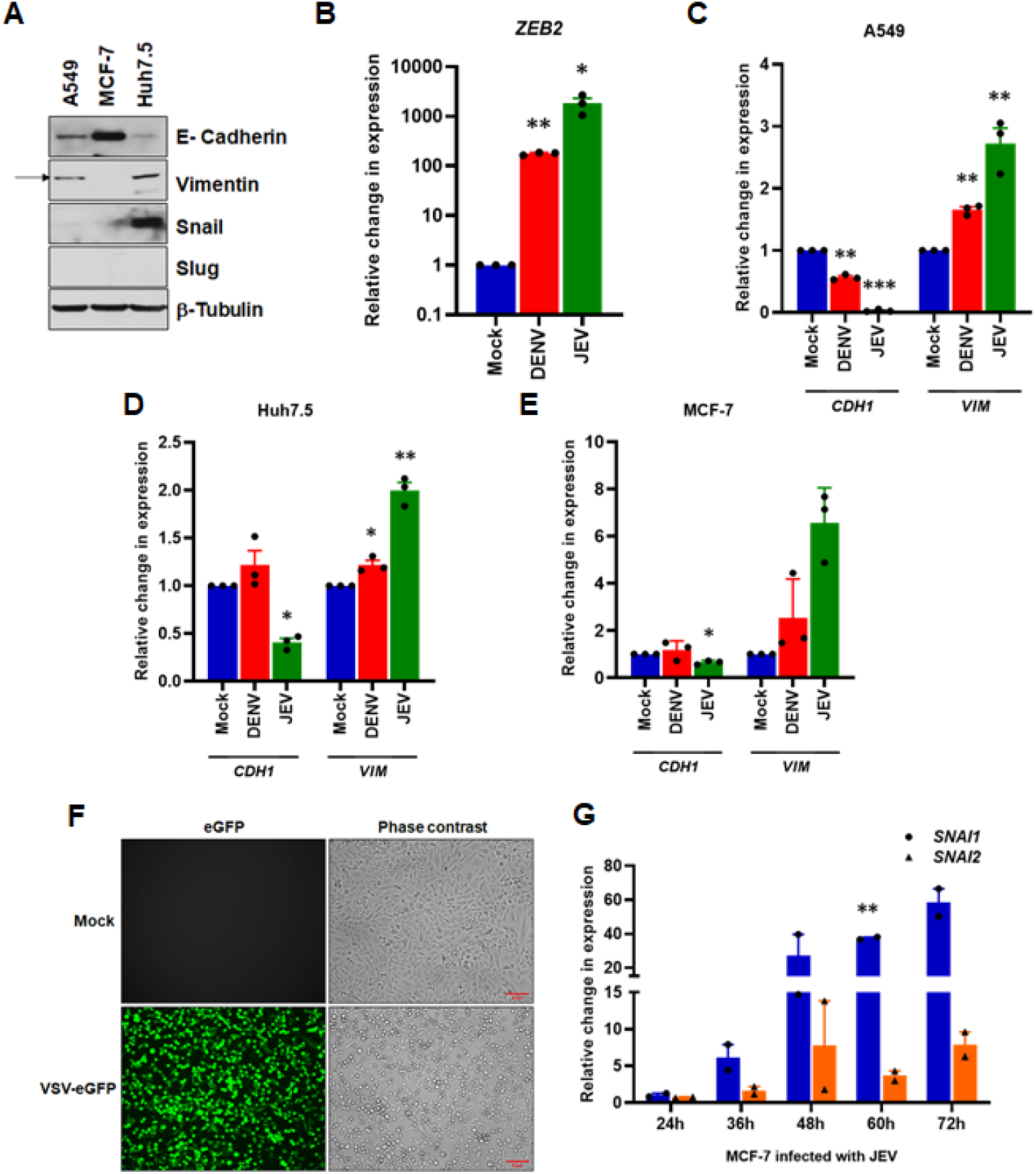
Non-oncoviruses induce EMT. (**A**) Profiling of EMT markers in three epithelial cancer cell lines A549, MCF-7, and Huh7.5 by immunoblotting. (**B**) Real-time qRT-PCR quantification of *ZEB2* in DENV and JEV infected A549 cells, as demonstrated in Figure 1B. (**C** - **E**) Real-time qRT-PCR analysis of DENV and JEV infected A549 (**C**), Huh7.5 (**D**), and MCF-7 cells (**E**), quantified for *CDH1* and *VIM* normalized against *GAPDH.* (**F**) Confirmation of VSV infection in A549 cells. A549 cells were infected with VSV-eGFP at 0.01MOI for 18 hrs and directly visualized using a fluorescence microscope. (**G**) Quantification of *SNAI1* and *SNAI2* transcripts in mock and JEV infected MCF-7 by real-time qRT-PCR at indicated time points. *GAPDH* was used as the normalization control.

**Figure S2:**
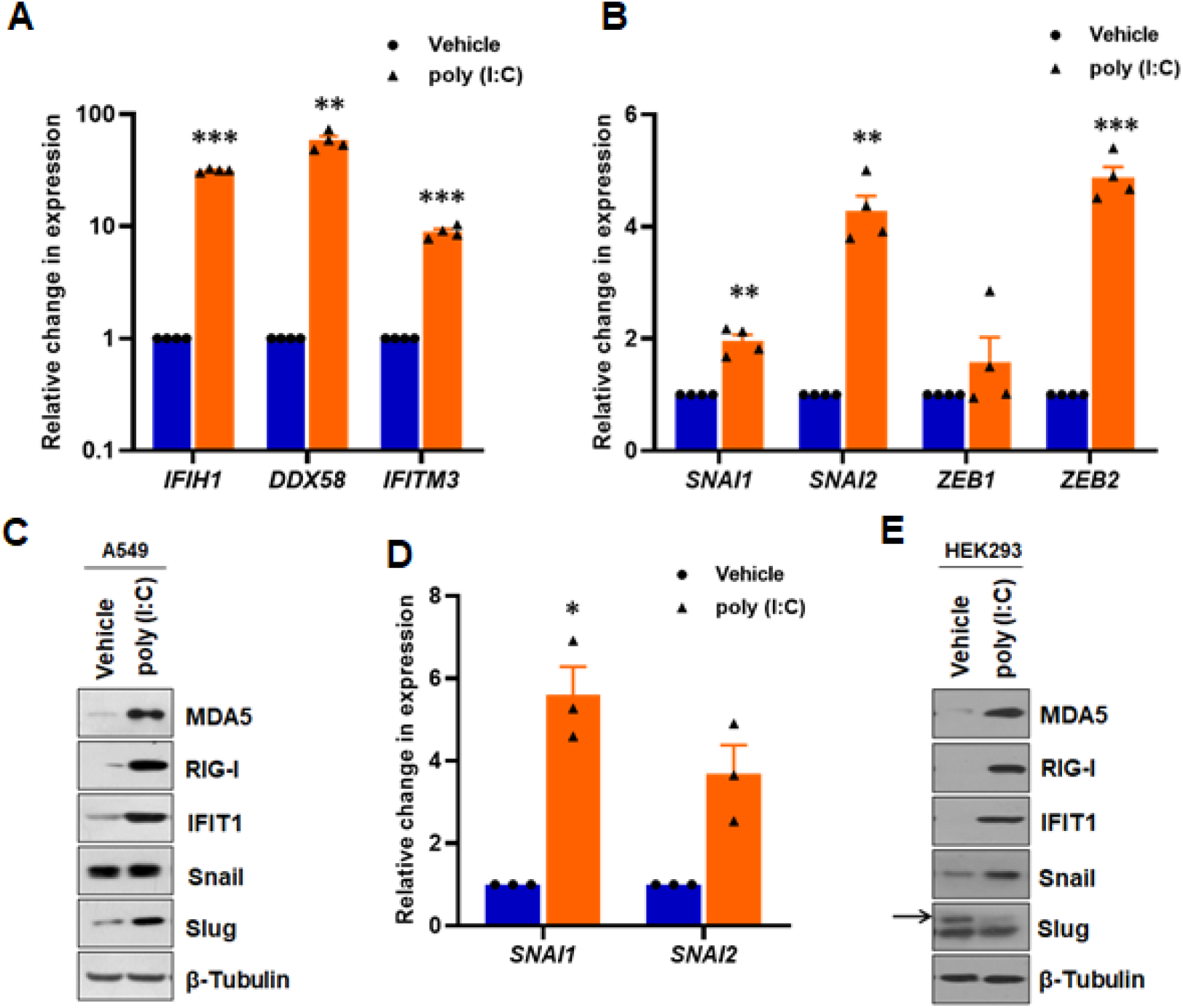
EMT-TFs are activated by viral RNA. Transfection of poly (I:C) into A549 cells (1000 ng/mL) and subsequent expression analysis of (**A**) antiviral genes, *IFIH1, DDX58,* and *IFITM3,* and (**B**) EMT-TFs, *SNAI1, SNAI2, ZEB1,* and *ZEB2,* by real-time qRT-PCR. *GAPDH* was used as the normalization control. (**C**) Expression levels of antiviral markers and EMT-TFs from vehicle and poly (I:C) transfected A549 detected by immunoblotting. (**D** & **E**) Analysis of EMT-TFs induction in HEK293 cells transfected with poly (I:C). Cells were transfected with 1000 ng/mL of poly (I:C) Quantitation of *SNAI1* and *SNAI2* (**D**) by real-time qRT-PCR in poly (I:C) transfected HEK293 cells. (**E**) Expression levels of antiviral markers and EMT-TFs from vehicle and poly (I:C) treated HEK293 detected by immunoblotting.

**Figure S3:**
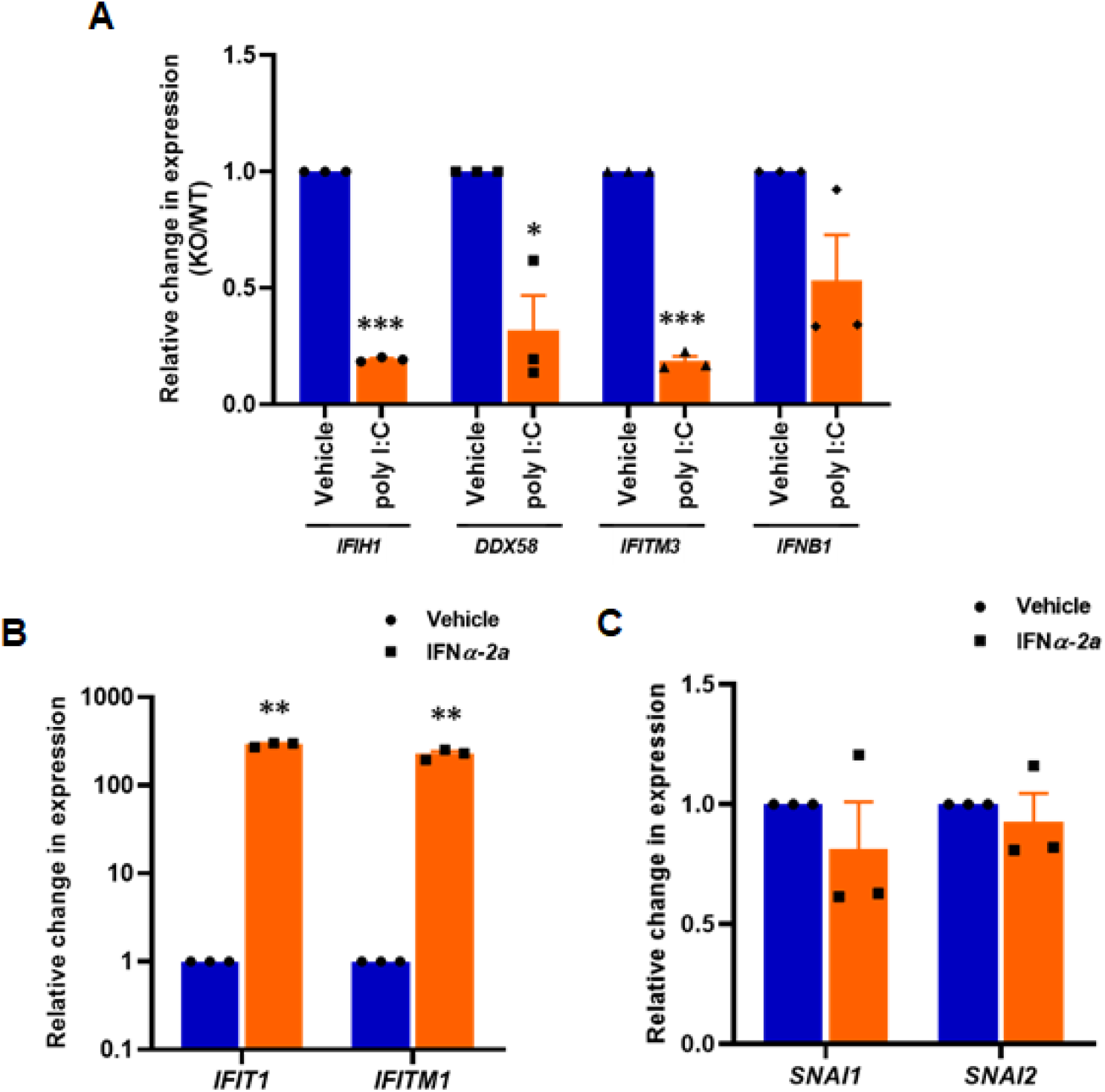
RLRs regulate EMT-TFs expression: (**A**) Effect of poly (I:C) treatment on antiviral genes, *IFIH1, DDX58, IFITM3,* and *IFNB1* in MAVS KO cells, compared to DWT cells. (**B and C**) Real-time qRT-PCR analysis to assess the effect of IFNα-2a on EMT-TFs treatment in MCF-7. Cells were grown to reach 70% confluency and treated with 50 ng/mL human IFNα-2a or PBS for 24 hrs. Cells were harvested and the respective transcripts were quantified.

**Figure S4:**
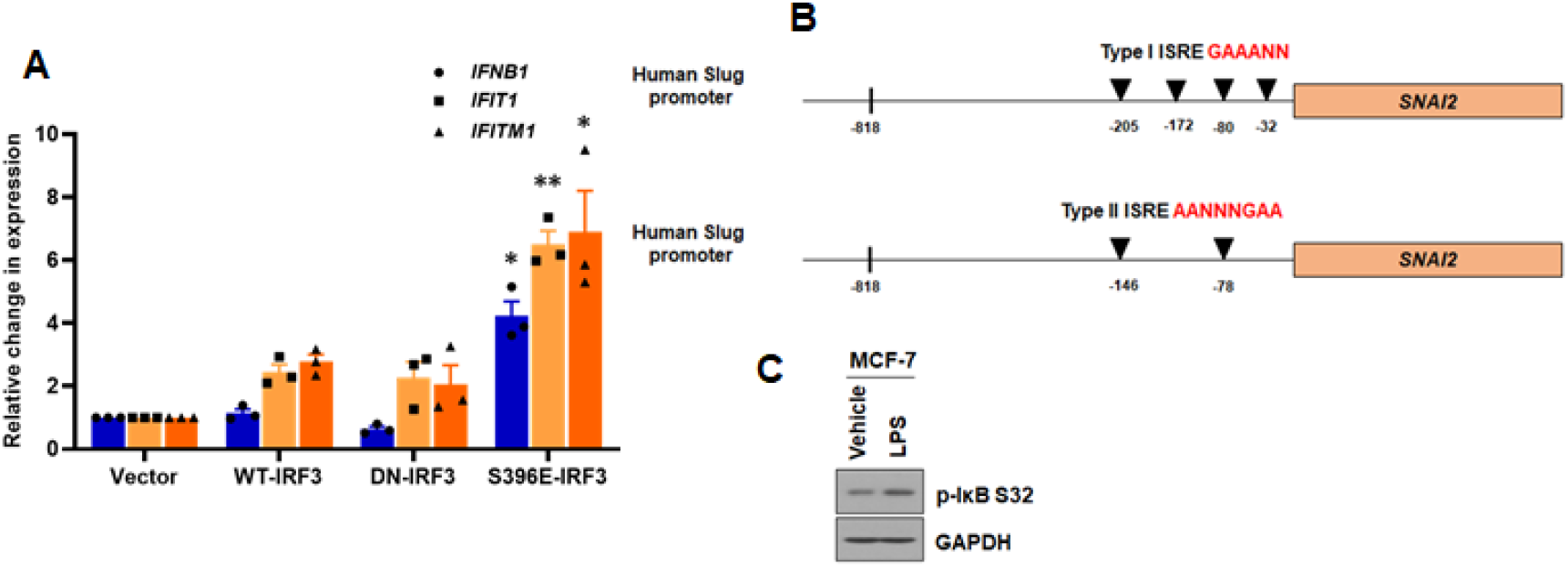
Phosphorylated IRF3 regulates *SNAI1* and *SNAI2* transcription: (**A**) Effect of over-expression of WT- and mutant IRF3 in MCF-7 cells on antiviral genes, *IFNB1, IFIT1* and, *IFITM1* upon quantified by real-time qRT-PCR. *GAPDH* was used as the normalization control. (**B**) Schematic representation of human Slug promoter for putative ISRE. Two types of ISRE, namely, type I – GAAANN and Type II – AANNNGAA, were identified in Slug promoter at indicated positions. (**C**) Immunoblot analysis of LPS treated MCF-7 cells for activation of the NF-κB pathway.

**Figure S5:**
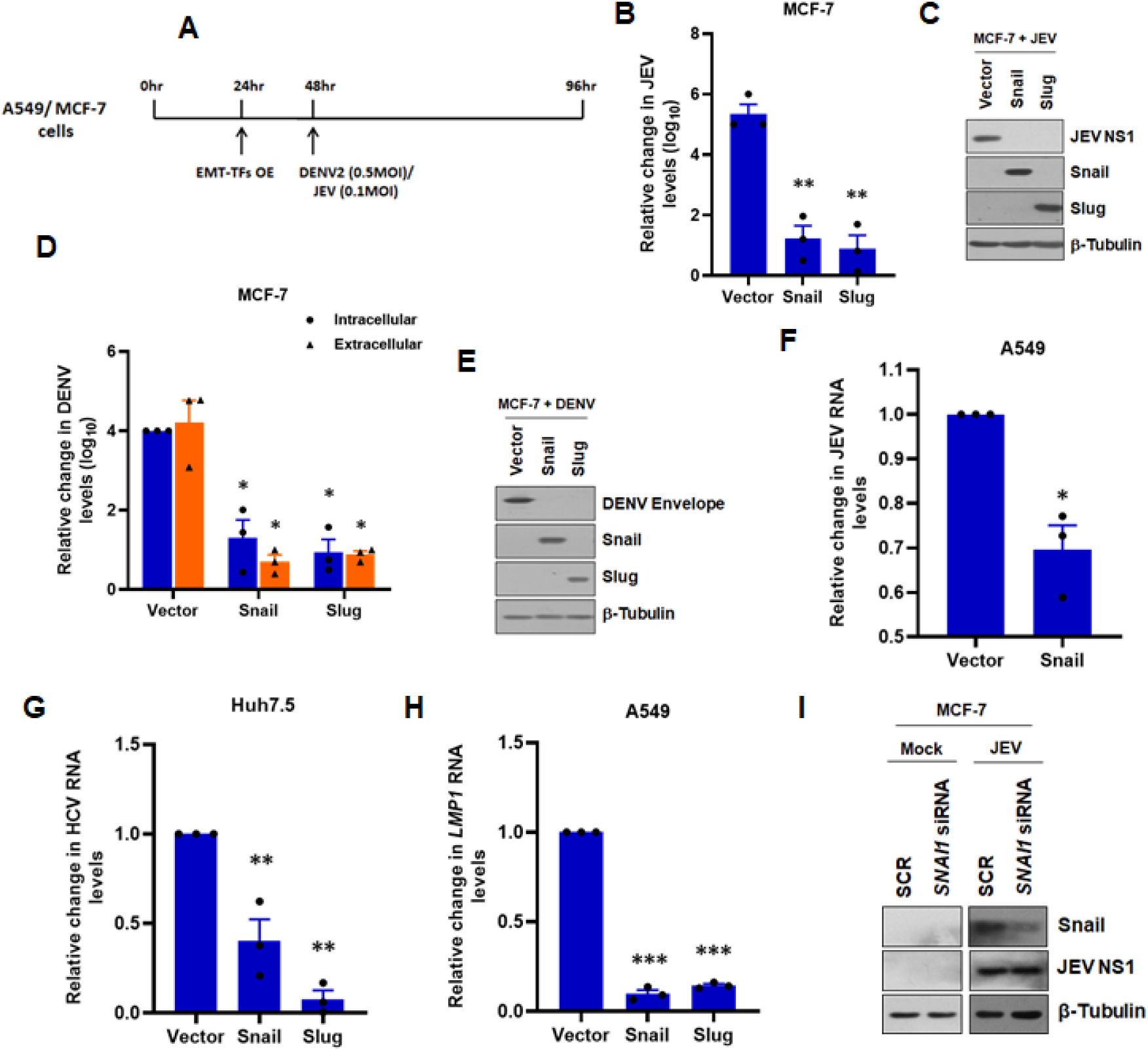
EMT-TFs suppress viral infections through its DNA binding activity: (**A**) Schematic representation of the experimental setup for panels 5**A** - 5**D**. (**B**) Snail/Slug over-expressing MCF-7 cells were infected with 0.1 MOI for 48 hrs. Cells were harvested and assessed for intracellular JEV titer by real-time qRT-PCR. Simultaneously, cells were lysed and immunoblotted (**C**) for JEV NS1, Snail, Slug, and β-Tubulin. (**D** & **E**) Similar to the panel (**B** & **C**), but DENV infection was done at 0.5 MOI for 48 hrs. Supernatant and cells were assayed for extracellular and intracellular titer (**D**) by foci forming assay and real-time qRT-PCR, respectively. (**E**) Immunoblot analysis for DENV envelope, Snail, Slug, and β-Tubulin. (**F**) For virus entry assay, A549 cells were transfected with Snail-expressing or empty vector for 24 hrs, followed by JEV infection at 1 MOI for 2 hrs. 2 hpi, cells were washed with PBS to remove unbound virus. Infected cells were treated with 50 μg/mL Proteinase K for 45 min to remove uninternalized virus, followed by treatment with PBS containing PMSF to inactivate Proteinase K. They were harvested and assayed for intracellular JEV RNA levels by real-time qRT-PCR. (**G**) Huh7.5 cells expressing Snail/Slug/vector were infected with HCV (0.5 MOI) for 72 hrs. Intracellular HCV RNA levels were measured by real-time qRT-PCR, and *GAPDH* was used as the normalization control. (H) Effect of Snail or Slug expression on EBV infection in A549 cells. 48 hpi cells were harvested, and intracellular *LMP1* RNA levels were quantified by real-time qRT-PCR. (I) MCF-7 cells were transfected with either 250 nM *SNAI1* siRNA or scramble using Lipofectamine 3000 for 24 hrs. Following transfection, cells were either mock or JEV infected at 0.1 MOI for 48 hrs. Cells were harvested and immunoblotted for Snail and JEV NS1 to confirm knock down and infection.

**Figure S6:**
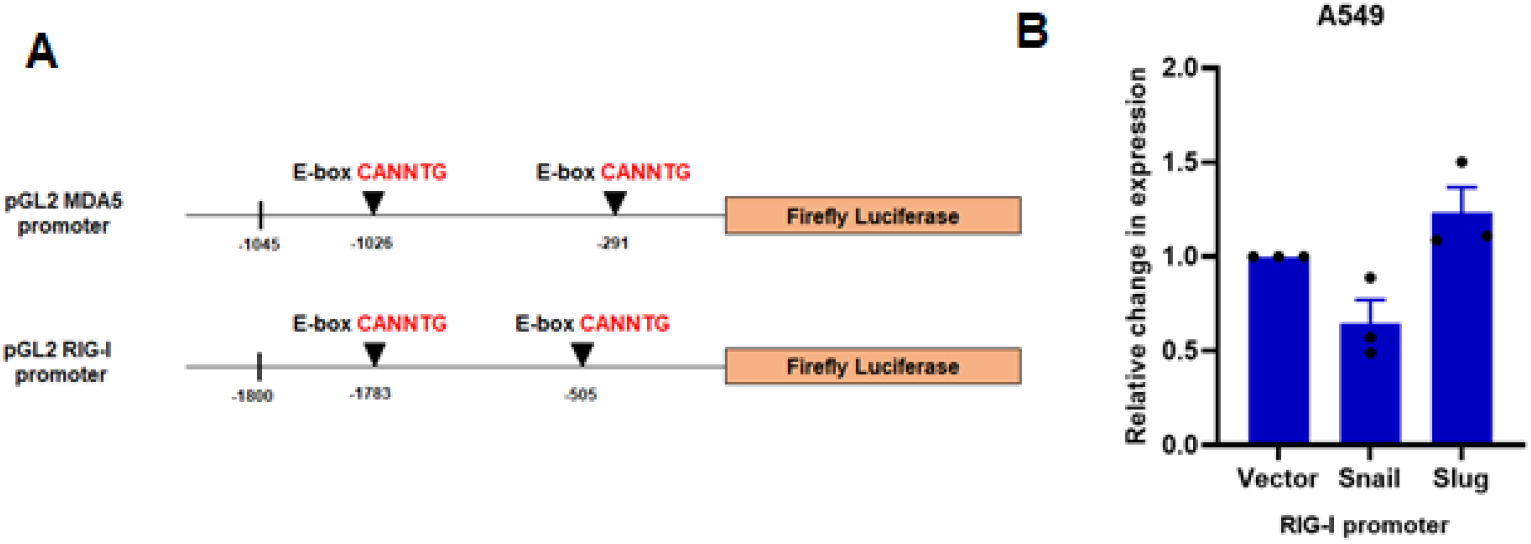
EMT-TFs enhance RIG-I levels, but not through transcription: (**A**) Schematic representation for canonical E-boxes located in *IFIH1* and *DDX58* promoter at mentioned positions. (**B**) Dual-luciferase assay in A549 cells co-transfected with pGL2 DDX58 promoter-reporter construct and Snail/Slug/vector expression constructs. pRL-CMV vector was used as the normalization control. Relative F/R ratios of luciferase activity are represented in the graph.

